# A new strategy for selection of crop cultivated varieties based on yield factors driving and meteorological prediction

**DOI:** 10.1101/766253

**Authors:** Rongsheng Zhu, Kai Sun, Xuehui Yan, Dawei Xin, Hongwei Jiang, Zhanguo Zhang, Yang Li, Zhenbang Hu, Zhaoming Qi, Chunming Yang, Qingshan Chen

## Abstract

Meteorological conditions, as an uncontrollable factor, have a direct impact on the growth and yield of crops in the field. It’s particularly important to choose the cultivated varieties suitable for environmental change. As for this problem, a new strategy for selection of crop cultivated varieties is proposed in this paper. The experiment selected Harbin pilot (Heilongjiang, China) and Changchun pilot (Jilin, China). In the experiment, the cultivated varieties suitable for forecasting meteorological conditions in 2015 could be provided according to the meteorological data of 2005-2014 years and the yield data of different soybean varieties in Harbin pilot. The same method was also taken in Changchun pilot. The results showed that the recommended varieties of Harbin pilot and Changchun pilot had 70% and 50% probability respectively for selection of soybean cultivated varieties when the yield evaluation trait was 100-seed-weight. As for seed weight per plant, the possibility of more than 60% of the recommended varieties in two pilots is to provide prejudgment. Therefore, this method can help to improve the current situation of the impact of uncontrollable meteorological conditions on crop yield and also can be applied to the selection of any other crops cultivated varieties under a certain environment.

## 1 Introduction

With the development of agricultural science and technology, some management factors such as pesticides, fertilizers and herbicides are basically stable and controllable, however, the effects of environmental factors such as precipitation, temperature and sunshine on crop growth are still in an uncontrollable state^1^. In recent decades, due to the changes of natural conditions and the impact of human activities, the global climate has shown a significant warming trend^2,3^, and agriculture is one of the areas that are sensitive to meteorological conditions^4^. Changes in meteorological conditions will have a significant impact on the ecological environment of agriculture as well as the growth and yield of crops^5^. Currently, many experts have done a lot of research on how meteorological conditions affect crop growth and yield^6-8^. For example, Chinese experts pointed out that low temperature treatment can slow the growth of soybean plants, delay the growth period, reduce the number of pods and the 100-seed-weight^9^. Mina and others simulated the influence of the future climate on the growth of maize by combining the climate regional model with the crop growth models such as WOFOST and CERES-Maize^10^. The key meteorological factors influencing spring soybean sowing to seedling, seedling and blooming period are the average daily temperature, and the key meteorological factor from emergence to maturity is precipitation, however, the key meteorological factors from flowering to maturity are sunshine hours and precipitation through the method of stepwise regression and correlation of path analysis^11^. Wang adopted APSIM model to analyze the effects of climate change on winter wheat phenology^12^. Pan described the increased demand for water as the plants grew faster. If the drought is encountered during the growth period, the number of shriveled grains and the 100-seed-weight will decrease, which will affect the yield of soybean^13^. Zhou and others introduced the impact of climate change on soil organic carbon and surface soil organic carbon storage and crop yield^14^. Chen and others analyzed the influence of meteorological factors on grain production in north China based on APSIM model^15^. Dong and others used the APSIM model to analyze the effect of drought on maize yield in northern China^16^.

According to the above research, it can be concluded that the environment has a direct impact on the growth and yield of crops. Research pointed out the biological significance of the genetic formula:phenotype = environment + genotype (P=E+G). The phenotype of an organism is the result of the accumulation and interaction of a particular genotype and the environment. But the genotype is built under the repeated effect of the environment, which is the genetic characteristic that the environment has given to the organism for a long time. It also confirms the idea that genotypes come from the environment^17^. Yan and He introduced that wheat starch traits were not only influenced by genotype and environmental accumulation factors, but also influenced by the interaction between genotype and environment^18^. Suppose the biological phenotype P is the yield in the formula, then yield will be affected by the internal genes of the crops and the external environment. Crop yields are generally affected by factors such as environmental 1 (meteorological factors), environment 2 (soil factors) and internal factors (genetic factors). Generally, it can be expressed as:

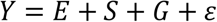

or

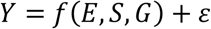

Where *E* is the environment, *S* is the soil, *G* is the gene.

According to the expression, the influence factor of yield can be more generally expressed as:

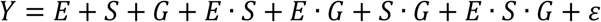

For the same place, *S* can be assumed to be constant, so *Y* = *f*(*E, G*) + *ε*. If it is assumed that the *ε* is contained in the environmental factor, then *Y* = *f*(*E, G*). However, both *E* and *G* are difficult to describe and the specific functional relationships are difficult to determine as well. Based on the above analysis, it is crucial to select suitable planting varieties according to meteorological conditions to improve crop yield. Therefore, if the weather conditions and crop cultivated varieties are known, an evaluation scheme for the yield will be given.

A new strategy for crop cultivated varieties selection based on meteorological predication is proposed to solve the problem of the influence of meteorological conditions on crop yield in this paper. The strategy adopts the weather prediction model based on phase space reconstruction^19^ and exponential smoothing^20^, and the evaluation classification model based on Euclidean distance^21^, which can achieve the optimal selection of planting varieties according to the target traits. This strategy provides a theoretical basis for the selection of crop cultivated varieties to improve crop yield.

In this paper, Harbin pilot (Heilongjiang Province, China) and Changchun pilot (Jilin Province, China) were selected as the test points. Harbin (N: 44°04′∼46°40′, E: 125°42′∼130°10′) and Changchun (N: 43°05′∼45°15′, E: 124° 18′∼127°02′)are important agricultural bases in China. The two pilots are high yield soybean, which belong to the temperate continental monsoon climate, with much land and few people, fertile soil and have a long history of planting soybean. However, in recent years, due to the changes of meteorological conditions, the market competitiveness of soybean in northeast China has decreased significantly. Most of the farmers are switching to other crops in the pursuit of higher returns, so the planting area of soybean significantly have been declining year by year.

In this experiment, the soybean varieties suitable for planting in the next year were selected according to the monthly meteorological data of Harbin from 2005 to 2014 and the meteorological data of Changchun from 2008 to 2009 through the use of the new selection strategy proposed in this paper. The experimental results showed that this strategy can predict the meteorological conditions of the next year accurately and can provide the recommended crop cultivated varieties through the classification evaluation model. This strategy also can help to improve the current situation of the impact of uncontrollable meteorological conditions on crop yield and it also has a certain significance for the planting of crops under a certain environment and condition.

## 2. Methods

### 2.1 Experiment materials

Compared with other meteorological conditions, precipitation, temperature and sunshine hours have a greater impact on soybean growth, which can be obtained by Shang^22^. During the growth cycle of soybean, the meteorological conditions of 5-9 months played an important role in the growth of soybean. Therefore, the meteorological data of monthly precipitation (Monthly precipitation refers to the depth of the liquid or solid that has been deposited from the sky to the surface of the ground in a certain month, without evaporation, infiltration, and loss of water.), monthly mean temperature (Monthly mean temperature refers to the average value of the average temperature of all days in a month.) and monthly sunshine hours (Monthly sunshine hours refers to the summation of the multi day time of the sun’s direct rays irradiating the ground within a month.) in Harbin during the past 2005-2015 years were selected as meteorological materials for Harbin pilot, which only included 5-9 months of meteorological data. The meteorological data from 2005 to 2014 are training samples, and the meteorological data of 2015 are testing samples. As for the Changchun pilot, the meteorological data of monthly precipitation, monthly mean temperature and monthly sunshine hours during the past 2008-2010 years were selected as meteorological materials, which also only included 5-9 months of meteorological data. The meteorological data from 2008 to 2009 are training samples, and the meteorological data of 2010 are testing samples. The meteorological data are provided by the statistical yearbook of Heilongjiang province and the statistical yearbook of Jilin province respectively^23^. The detailed meteorological data refer to the Supplementary Table S1-S6 online.

The experiment of Harbin pilot was conducted in the experimental field of Northeast agricultural university from 2005 to 2015. A total of 149 soybean germplasm resources were used as the experiment materials, which are provided by Northeast Agricultural University, Harbin, Heilongjiang, China and cover the main varieties at home and abroad.

The experiment of Changchun pilot was conducted in the experimental field of Jilin academy of agricultural sciences from 2008 to 2010, and 274 soybean germplasm resources were selected as the experiment materials, which are provided by the Agricultural Biotechnology Research institute, Jilin Academy of Agricultural Sciences (JAAS). Detailed varieties and phenotype data are shown in Supplemental Table S7 and Table S8 online.

The planting time was at the end of April every year, and the harvest time was in late September every year. The experiment was conducted in a randomized block group with three rows in each group, and three repeated tests were required in the process of planting. The row distance was 50cm, the total length of the row was 300cm, and the plant spacing of soybean in the two pilots was 6±1cm. The seedling, weeding and strengthening of the cultivated soil were carried out at seedling stage. In the process of flowering pod, the pod should be prevented from falling off. In the late flowering period, the largest soybean leaf area was guaranteed to be photosynthetic, and the water was replenished in time when the bean grain was about to mature.

### 2.2 Experiment model

The main experimental model of this paper is the selection model of crop cultivated varieties with single target and single location, such as the 100-seed-weight, seed weight per plant, seed number per plant, seed number per pod, plant height and so on, which can be used as the yield target traits. Plant varieties can be selected according to the requirements of planting by selecting models of crop cultivated varieties.

The crop cultivated varieties selection model for single location and single target trait is as follows:

Suppose:

*y*_1_, *y*_2_ … *y*_*n*_ is the target traits of *n* years *m* species (such as 100-seed-weight or other traits)
*p*_1_, *p*_2_ … *p*_*n*_ is the monthly precipitation of *n* years (mm)
*t*_1_,*t*_2_ … *t*_*n*_ is the monthly mean temperature of *n* years (°C)
*s*_1_,*s*_2_ … *s*_*n*_ is the number of sunshine hours of n years (h)

The specific steps of the algorithm are as follows:

**Step 1:** Predict the average temperature *t*_*n*+1_, monthly precipitation *r*_*n*+1_and monthly sunshine hours *s*_*n*+1_of the *n* + 1 year.

The meteorological conditions of the *n* + 1 year can be obtained based on the prediction model of certain time series, and the results are as follows.

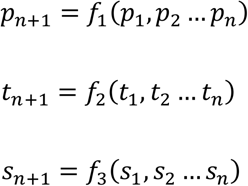

Therefore, we can obtain the meteorological time series:

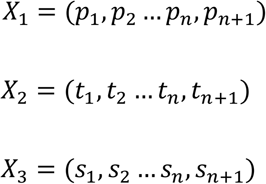

**Step 2:** Standardize meteorological time series.

According to the formula below

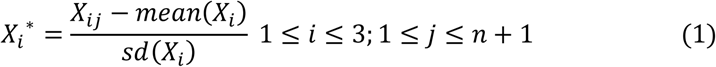

The results of the standardized time series are as follows:

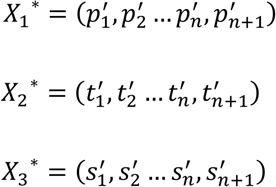

**Step 3:** Select a distance (usually the Euclidean distance) as a criterion. The Euclidean distance between the meteorological conditions of the*n* + 1year and the meteorological conditions of the previous *n* years is calculated (equation (2)).

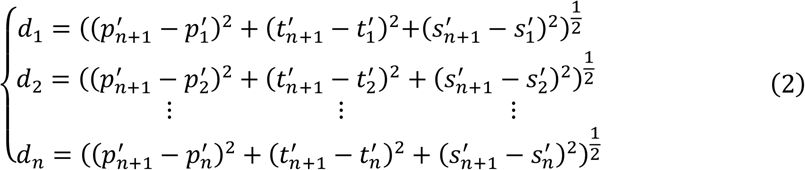

**Step 4:** Order *d*_1_, *d*_2_ … *d*_*n*_ from small to large scale.

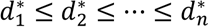

and then the *Y*_*k*_ (1 ≤ *k* ≤ *n*)year corresponding to the shortest Euclidean distance 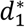 is selected.

**Step 5:** Sort crop cultivated varieties of *Y*_*k*_ (1 ≤ *k* ≤ *n*) year in accordance with the yield target traits. Threshold value α is set according to the need of planting, and the top α% varieties of the *Y*_*k*_ year are selected to form the recommended varieties set *P* for the *n* + 1 year.

#### 2.2.1 Meteorological prediction model based on phase space reconstruction

The theory of phase space reconstruction proposed by Packard in 1980, and the theory holds that the evolution of any component in the system is determined by the other components that interact with it. Therefore, the information of these relevant components is implied in the development of any component.

For a nonlinear system, the observed value is recorded as time series {*x*(*t*_*i*_),*i* = 1,2, …, *n*)}, and the observed value can be used to form the *m* dimension vector:

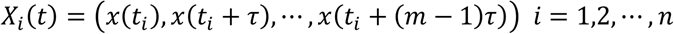

Where *τ* is a delay time, *m* is embedded dimension, and *X*_*i*_(*t*) is a phase point in the constructed space.

*Takens* proves that the phase space is reconstructed by proper embedding dimension *m* and delay time *τ*. The “trajectory” in the embedded space is equivalent to the original system^19^. *d* ≥ 2*n* + 1 is a sufficient condition for reconstructing the embedding dimension of phase space, and *n* is the number of independent parameters in the dynamic system. But Jiang points out that the necessary conditions have not been documented in the literature^24^. Therefore,

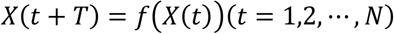

Here, *T*(*T* > 0) is the prediction model for forward prediction step, and *f*(·) is the prediction model of reconstruction.

According to the attractor in the fitting phase space, it can be divided into two kinds: the global method and the local method. The global method can get the *f*(·) by regarding all the points in the trajectory as the fitting objects. Thus, it is feasible in theory to predict the trend of trajectory. However, it is difficult to predict accurately when the phase space structure is more complex. The local method is to take the last point of the phase space trajectory as the center point and the recent locus of the center point as the relevant point, and then the relevant points are proposed to be combined to estimate the direction of the next point. Finally, the predicted values are separated from the coordinates of the predicted trajectories. Compared with the two methods, the application of local method is more extensive^25^.

#### 2.2.2 Exponential smoothing prediction model

The exponential smoothing method, proposed by American Brown in 1959, is one of the most commonly used methods for analysis of time series, which is mainly applied to the short-term forecast and is a kind of excellent performance, strong adaptability of prediction methods^26^. This method assumes that the predicted value of the future is related to the known data, and the recent data has a large impact on the predicted value, but the long-term data has less influence on the predicted value. Therefore, the influence shows the trend of decreasing geometrical series. This method is widely used in various fields^27-29^.

The exponential smoothing model consists of an exponential smoothing method, the double exponential smoothing and the cubic exponential smoothing method. The basic formula (equation (3)) of exponential smoothing is:

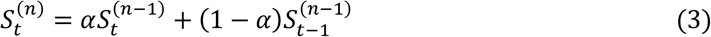

Where *S*_*t*_ is the exponential smooth value of the *t* cycle.*α*(0 ≤ *α* ≤ 1) is a smooth exponent. *t* is a periodic number.

The simple exponential smoothing method is suitable for the time series analysis with horizontal trend, which can only predict the time series of the short term. If the time series has an upward or downward trend, it is best to predict with the double exponential smoothing. If the time series not only has the obvious rise or decrease trend but also presents the seasonal change, then the cubic exponential smoothing method is selected.

##### (1) The selection of initial value

In general, the observation value of the first phase can be selected as the initial value when the time series data items are large. If the number of the original numbers is relatively small, the average number of the first several periods (Usually the first three data) can be selected as the initial value.

##### (2) The selection of smoothness index σ

The smoothness index σ is a proportional factor for the different effects of the new and old data in the prediction, which not only represents the reaction speed of the model to time series data, but also determines the ability of prediction model to repair the error. The larger the value of σ, the greater the impact of the recent observed values, and the more dependent on the recent data, the higher the sensitivity of the model. Conversely, the smaller σ is, the slower the model changes, and the prediction results are more dependent on the early data.

There are three ways to determine the smoothness index σ :Difference-ratio-mean method, Experiential judgment method and Trial calculation method. The general selection criteria are as follows:

1. When the observed values show a stable horizontal trend, σ is generally selected between 0.1-0.3. The weight of each period is not very different, which reduces the magnitude of correction. Meanwhile, the new sequence contains more original sequence information, which helps to increase the credibility of prediction results.
2. When the observed values fluctuate greatly, σ is generally selected between 0.3-0.5.
3. When the observed value fluctuates greatly and presents obvious seasonal variation, the value range of σ is 0.6-0.8, so as to enhance the role of the recent observed value in the prediction process^30^.

In this paper, σ is selected by Trial calculation method, and the determination of σ is generally evaluated by the minimum *RMSE* (root-mean-square error) (equation (4)) of the predicted results, and then the most reasonable σ value is selected^31^.

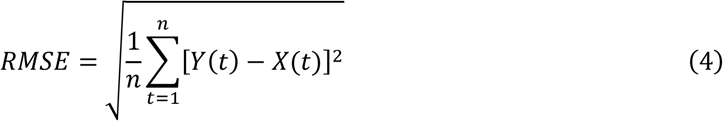

Where *n* is the number of predicted data; *Y*(*t*) represents the predicted values of the *t* period; *X*(*t*) is the actual value of *t* period.

#### 2.1.3 Evaluation classification model of crop cultivated varieties based on Euclidean distance

Suppose that the monthly precipitation, the monthly mean temperature and the monthly sunshine hours matrix of *n* years are respectively A, B and C.

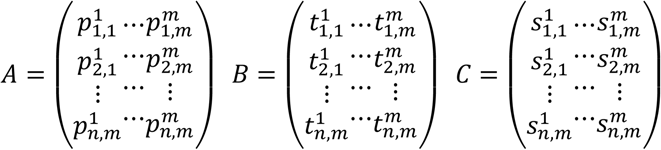

In the matrix, *m* is the month of the observed value. According to the meteorological data of the previous *n* years, the prediction data matrix of the monthly precipitation, monthly average temperature and monthly sunshine hours of *n* + 1 year can be obtained by using phase space reconstruction and exponential smoothing prediction model. Then a multi-dimensional matrix *D* with meteorological conditions of observed and predicted values can be obtained.

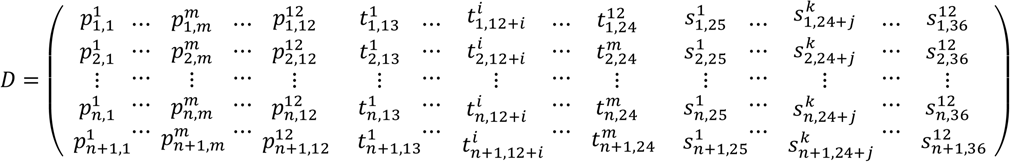

Where *m*(1 ≤ *m* ≤ 12) is the month of precipitation, *i*(1 ≤ *i* ≤ 12)is the month of average temperature, and *k*(1 ≤ *k* ≤ 12) is the month of sunshine hours.

The steps of the Euclidean distance evaluation classification model are as follows:

**Step 1:** Matrix *D*′ can be gained after standardizing the column vectors of the multi-dimensional matrix *D* according to Eq. (1).

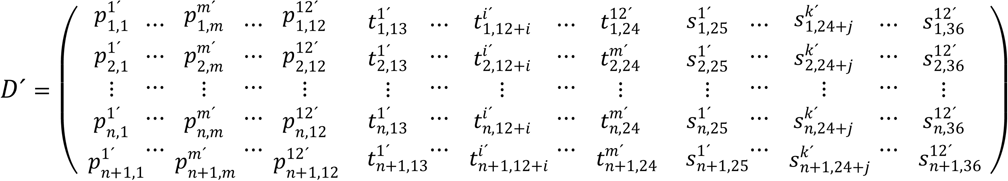

**Step 2:** Calculate the Euclidean distance between the *k* rows (*k* = 1,2 … *n*), and the *n* + 1 row respectively by equation (5).

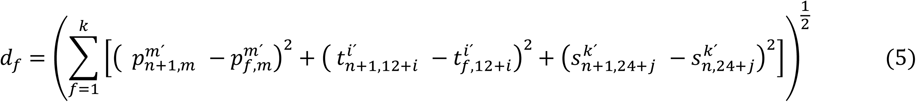

**Step 3:** Order *d*_1_,*d*_2_ … *d*_*n*_ from small to large scale.

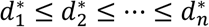

**Step 4:** Select the year *Y*_*k*_ (1 ≤ *k* ≤ *n*)that corresponds to the shortest Euclidean distance. In the *Y*_*k*_ (1 ≤ *k* ≤ *n*) year, the actual yield of the crop is sorted according to the target traits, and then the suitable threshold α is selected to classify and select the varieties according to the actual needs.

### 2.3 Experiment methods

#### 2.3.1 Meteorological prediction method based on phase space reconstruction

In the prediction experiment of precipitation and sunshine, the experimental data materials of the monthly precipitation and monthly sunshine hours of Harbin pilot were converted into line graphs, and the results were shown in Fig. 1. The data line chart of monthly precipitation and monthly sunshine hours in Changchun pilot

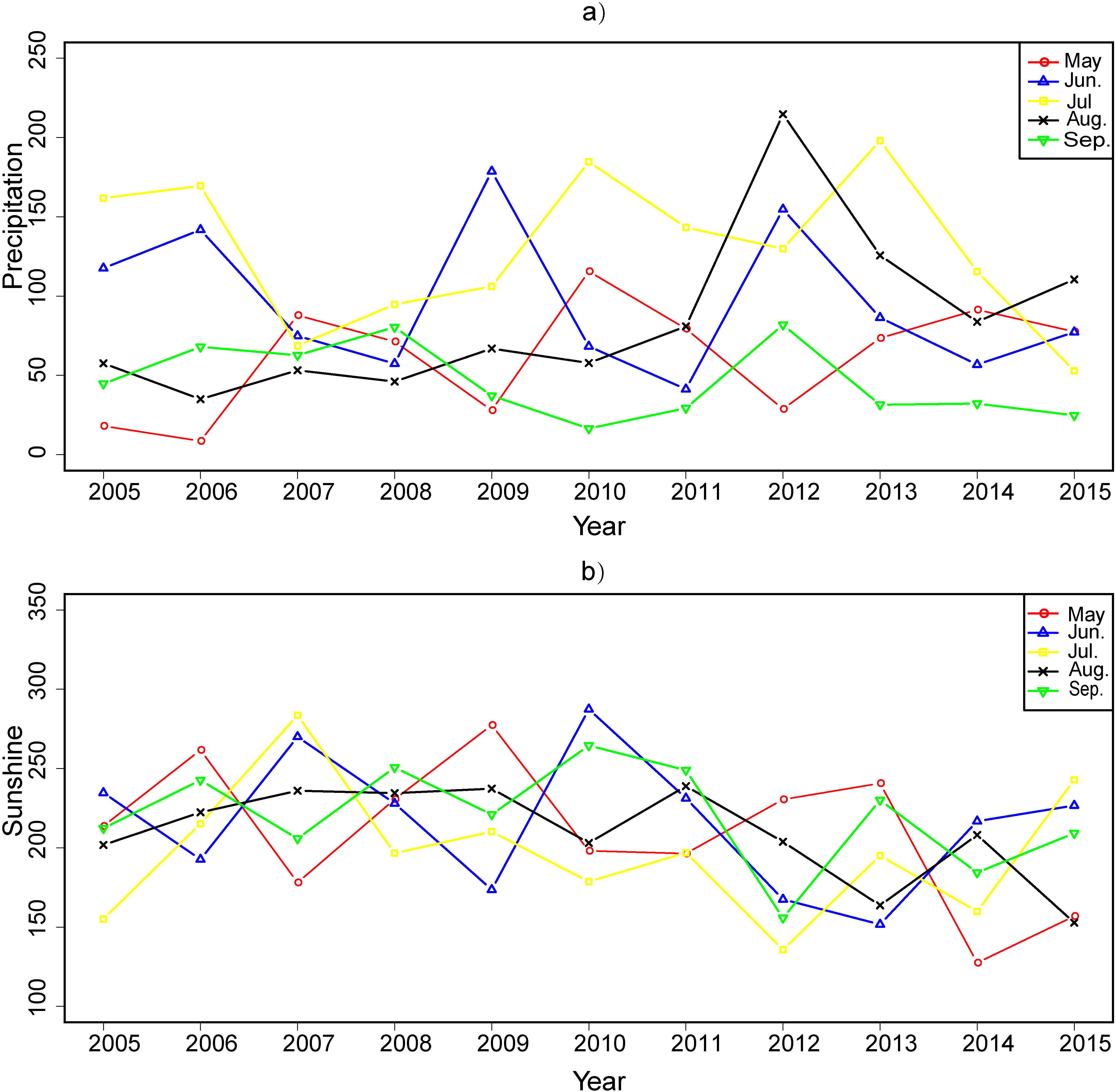

Fig. 1 shows that the data of the monthly precipitation and the monthly sunshine hours in Harbin pilot show an obvious nonlinear trend. Therefore, the prediction model based on phase space reconstruction were chosen to predict precipitation and sunshine hours, the concrete steps are as follows:

**Step 1:** The time series *x*(*n*), *n* = 1,2, …, *N* can be obtained after zero mean processing according to the data of monthly precipitation and sunshine hours in the period from 2005 to 2014.

**Step 2:** Determine phase space saturation embedding dimension *m* and time delay *τ*. A self-correlation method was used to determine the time delay in phase space reconstruction^32,33^. The commonly used methods for determining the best embedding dimension *m* were pseudo nearest neighbor^34^, G-P method^35^, singular value decomposition method^36^, etc. In addition, there were various improved algorithms, such as C-C^37^ and G-Z algorithms^38^, etc. To sum up, pseudo nearest neighbor point method is a method with more application. The maximum Lyapunov exponent of the chaotic time series is calculated by using small-data method to prove that the time series of the monthly precipitation and the monthly sunshine time series have chaotic characteristics^39^.

According to time delay *τ* and embedding dimension *m*, time series *x*(*n*), *n* = 1,2, …, *N* is converted into the following results.

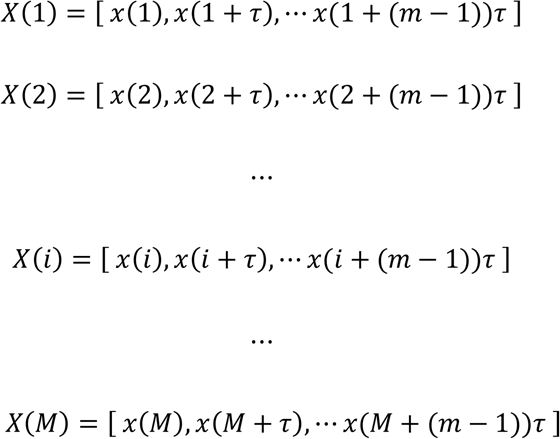

Where *M* is the space point for reconstructing space, *M* = *N −* (*m −* 1)*τ*.

Suppose: *Y*(*m,i*) = *X*(*i*) = [*x*(*i*),*x*(*i* + *τ*), … *x*(*i* + (*m* − 1))*τ*]

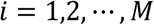

Therefore, {*Y*(*m,i*), *i* = 1,2, …, *M*} is the subspace of *R*^*m*^ space, which is the phase space required for the experiment.

**Step 3:** Search for near neighbors. In the phase space, suppose the current state point is *X*(*j*), and the neighborhood point of *X*(*j*) is found by calculating the Euclidean distance between the points *X*(*i*) *i* = 1,2, …, *M* and the *X*(*j*).

**Step 4:** Find the next phase point *X*(*i*_1_ + 1), *X*(*i*_2_ + 1), …, *X*(*i*_*p*_ + 1)for all the nearest neighbors. The coefficient matrix *A* and *B* are obtained by fitting the space trajectory in this small neighborhood.

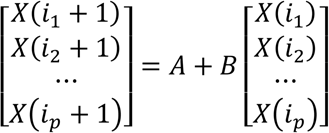

After obtaining *A* and *B*, the trend of trajectories in phase space can be get according to formula *X*(*j* + 1) = *A* + *BX*(*j*), so the predicted value of *x*(*j* + 1 + (*m −* 1)*τ*)in monthly precipitation and monthly sunshine hours is obtained.

#### 2.3.2 Exponential smoothing weather prediction method

In the prediction experiment of average temperature, the average temperature test material data of Harbin pilot was converted into a line chart, as shown in Fig. 2. The line chart of the monthly mean temperature test materials for Changchun pilot is shown in Supplemental Fig. S2 online. Fig. 2 shows that the time series data of the monthly mean temperature show a relatively stable horizontal trend, so an exponential smoothing method was chosen to predict.

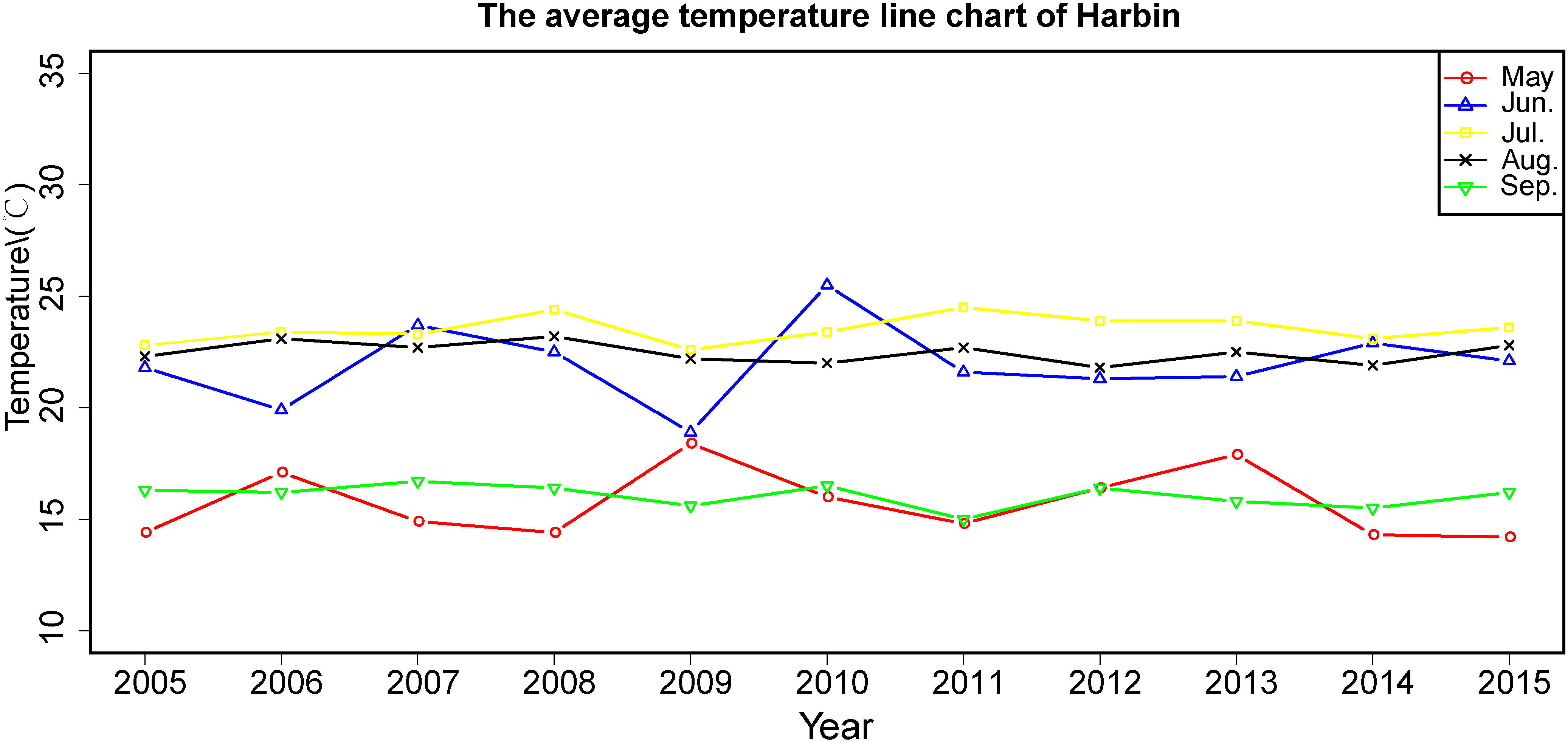

An exponential smoothing prediction method is as follows:

**Step 1:** An exponential smoothing prediction method formula.

It is assumed that the observed values of the time series for the 5-9 months period of 2005-2014 years are *x*_*i*_(*i* = 1,2 …). The exponential smoothing value of phase *n* is 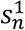, and the corresponding formula is equation (6):

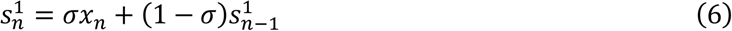

Where *x*_*n*_ is the actual value of phase *n*. σ is the smoothness index; 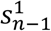 is the smooth value of the previous period; 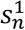 is the predicted value of the *n* + 1 period, which is the smoothing value of this period.

**Step 2:** The selection of initial value.

The number of the observed value of Harbin pilot is 10, which is less than 15. Therefore, the average value of the first three data of the observed values is used as the initial value of the smoothing prediction (equation (7)).

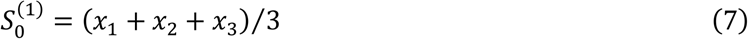

**Step 3:** The selection of smoothness index σ.

According to Fig 2, it can be concluded that the observed values show a relatively stable horizontal trend, so the smoothness index should be smaller, generally between 0.1 and 0.3. In this paper, 0.1, 0.2 and 0.3 were selected to predict the average monthly temperature respectively, and at the end of the prediction, the *σ* value corresponding to the minimum *RMSE* can be used as the smoothness index of the monthly average temperature prediction.

#### 2.3.3 Evaluation classification method based on Euclidean distance

Multi-dimensional matrix *D* can be obtained according to Euclidean distance evaluation classification model and meteorological materials. In this paper, Harbin pilot is taken as an example. Therefore, the first ten lines in the multi-dimensional matrix *D* represent the actual meteorological conditions of 2005-2014. The eleventh line stands for the forecast weather conditions for 2015. *m, i* and *k* range from 5-9, so the matrix *D* can be expressed as:

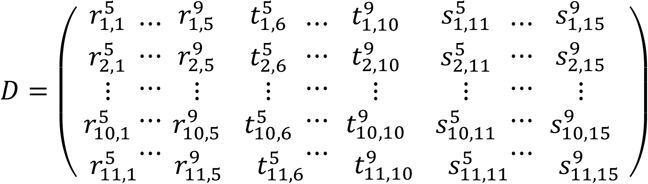

**Step 1:** Matrix *D*′ can be gained after standardizing the column vectors of the multi-dimensional matrix *D*.

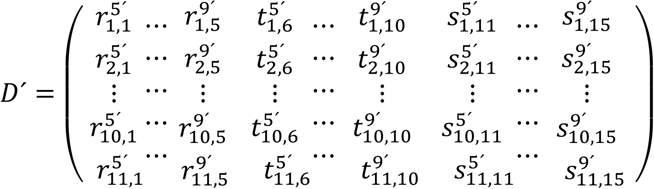

**Step 2:** The Euclidean distance between the first ten lines of the matrix *D*’ and the eleventh line of the matrix is calculated by equation (5). Then, according to the steps 3 and 4 in the evaluation classification model, the classification selection of the varieties of Harbin pilot was completed. The calculation method of the Changchun pilot was the same.

## 3 Results

### 3.1 The Predicted results of meteorological conditions

In the prediction experiment of monthly precipitation, the time delay *τ* and the embedding dimension *m* were calculated by self-correlation method and the pseudo nearest neighbor point method in Harbin pilot, so the time delay *τ* = 2 and the embedding dimension *m* = 4. The maximum Lyapunov index of the chaotic time series was calculated by small-data method, which was 1.6234 and indicated that the monthly precipitation time series was chaotic. The same method was used in Changchun pilot to obtain the time delay *τ*, the embedding dimension *m* and the maximum Lyapunov exponent, and the results were 2, 4 and 1.3728 respectively. In the experiment of monthly sunshine hours, the time delay *τ* and embedding dimension *m* of Harbin pilot and Changchun pilot were obtained by the self-correlation method and the pseudo nearest neighbor point method respectively. The results were that the time delay *τ* = 2 and the embedding dimension *m* = 6 in Harbin pilot, the time delay *τ* = 2 and the embedded dimension *m* = 4 in Changchun pilot.

A prediction model based on an exponential smoothing method was used in the prediction experiment of monthly mean temperature. According to the distribution trend of monthly mean temperature data and the selection criteria of sigma, the smoothness index 0.1, 0.2 and 0.3 were selected for meteorological prediction. The *RMSE* values corresponding to different smoothness index were shown in Table 1 and Fig. 3. The *RMSE* results of Changchun pilot were shown in Supplemental Fig. 3 online. Therefore, according to Fig. 3, the smoothness index corresponding to the minimum *RMSE* value of May to September was selected for meteorological prediction. Therefore, in the experiment of the average temperature forecast from May to September in 2015 in Harbin pilot, the smoothness index was 0.3, 0.2, 0.3, 0.2 and 0.3 respectively (Fig. 4). In Changchun pilot, the average temperature of May to September in 2010 was predicted by the smoothness index 0.3, 0.2, 0.3, 0.2 and 0.3(Supplemental Fig. 4 online). The prediction meteorological data of the Harbin pilot in 2015 and the Changchun pilot in 2010 were obtained through the above methods (Table 2). As shown in Table 2, the meteorological prediction model can accurately predict the meteorological conditions in the coming year and provide theoretical data support for the implementation of the strategy.

**Table 1.**
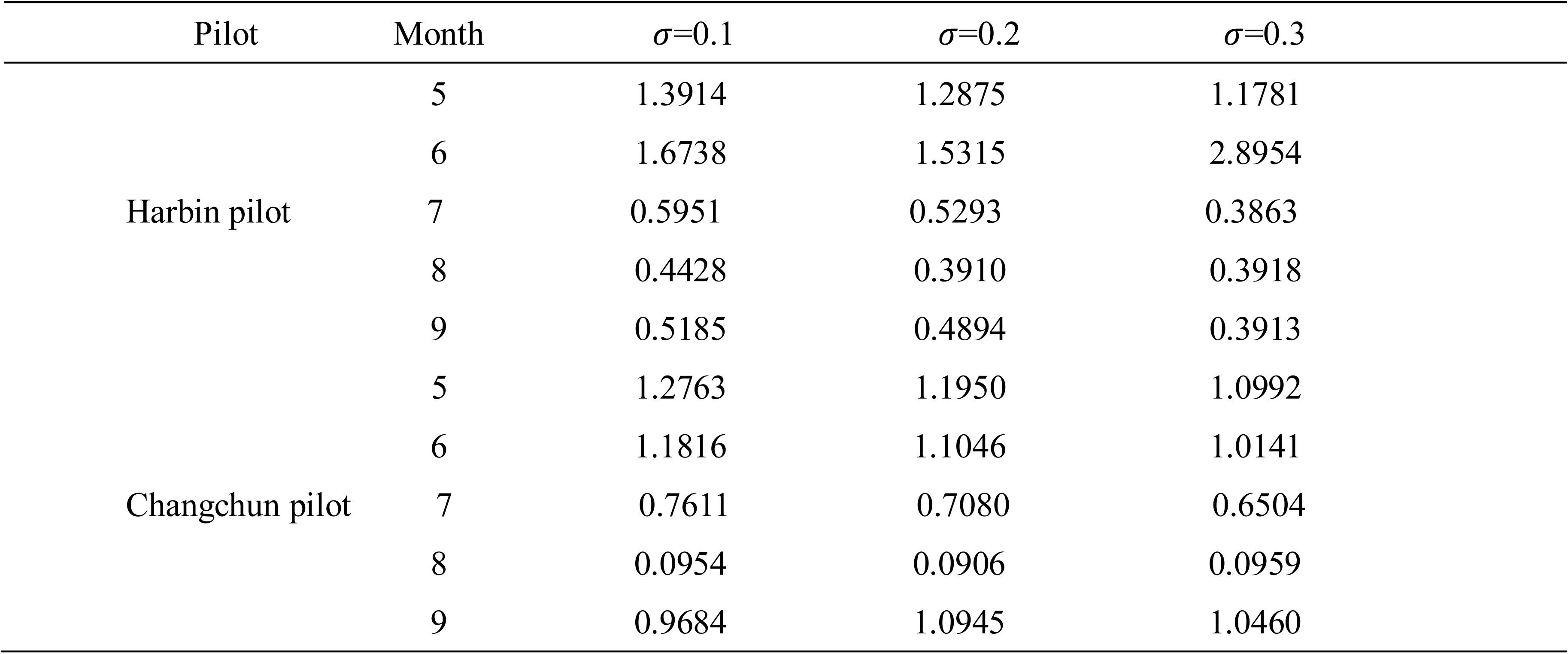
The *MSE* value of different smoothness exponent *σ* in the monthly mean temperature prediction experiment of Harbin pilot and Changchun pilot in May to September.

**Table 2.**
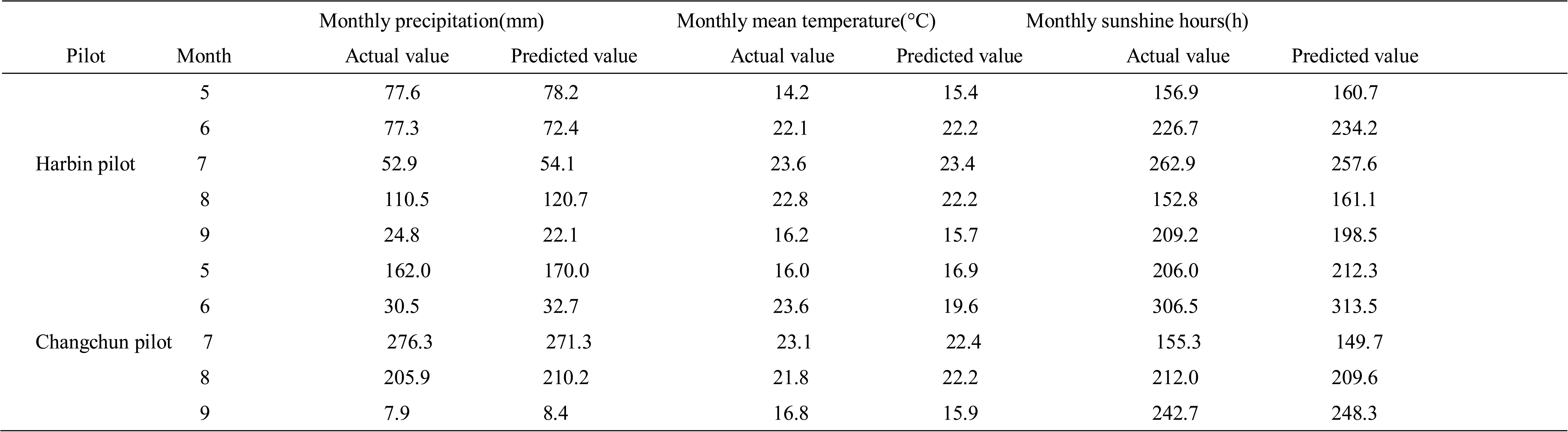
The forecast data of two pilot meteorological conditions are compared with the actual data. The pilot data of Harbin is the actual meteorological data and forecast meteorological data for 2015. The pilot data of Changchun is the actual meteorological data and forecast meteorological data for 2010.

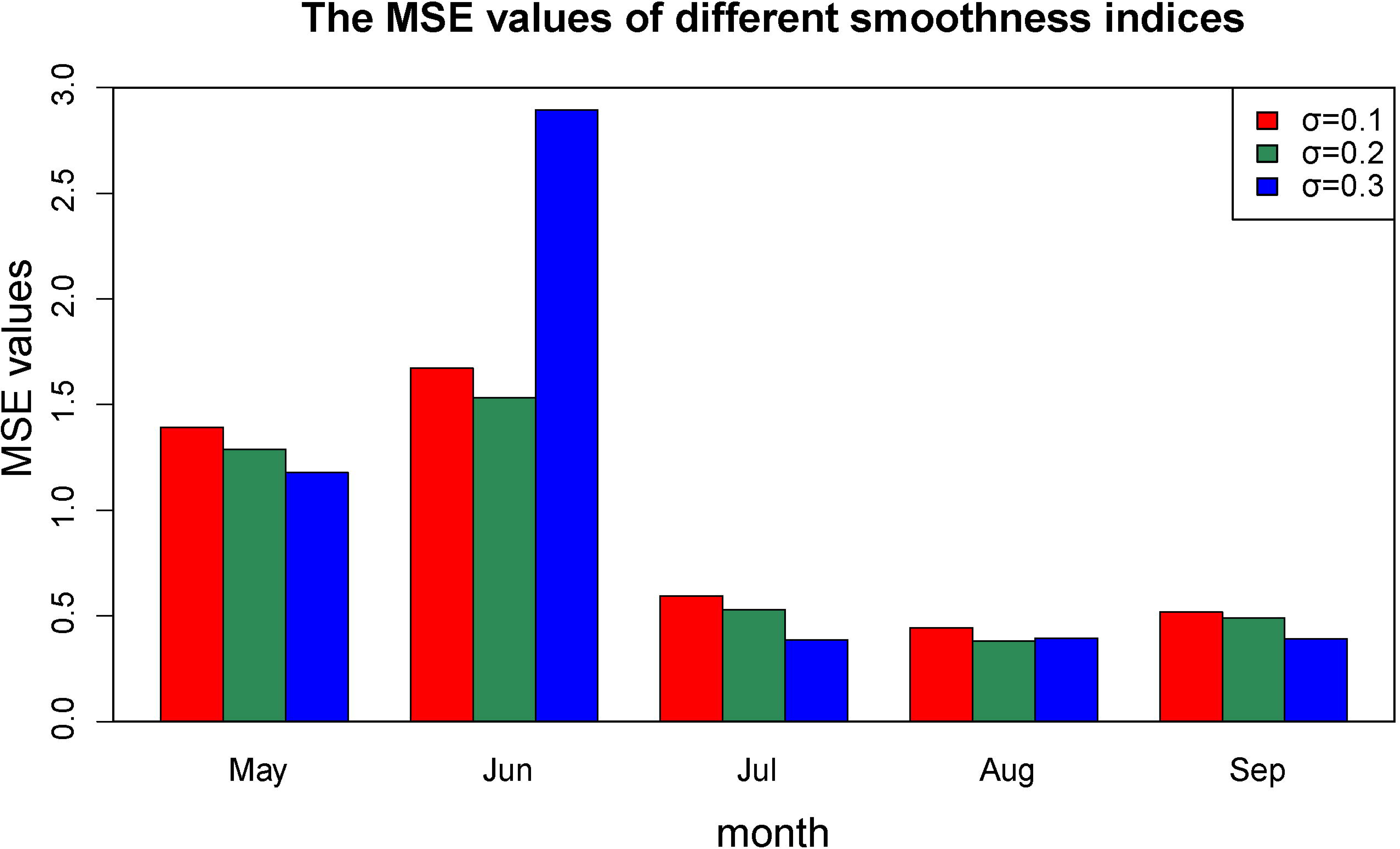

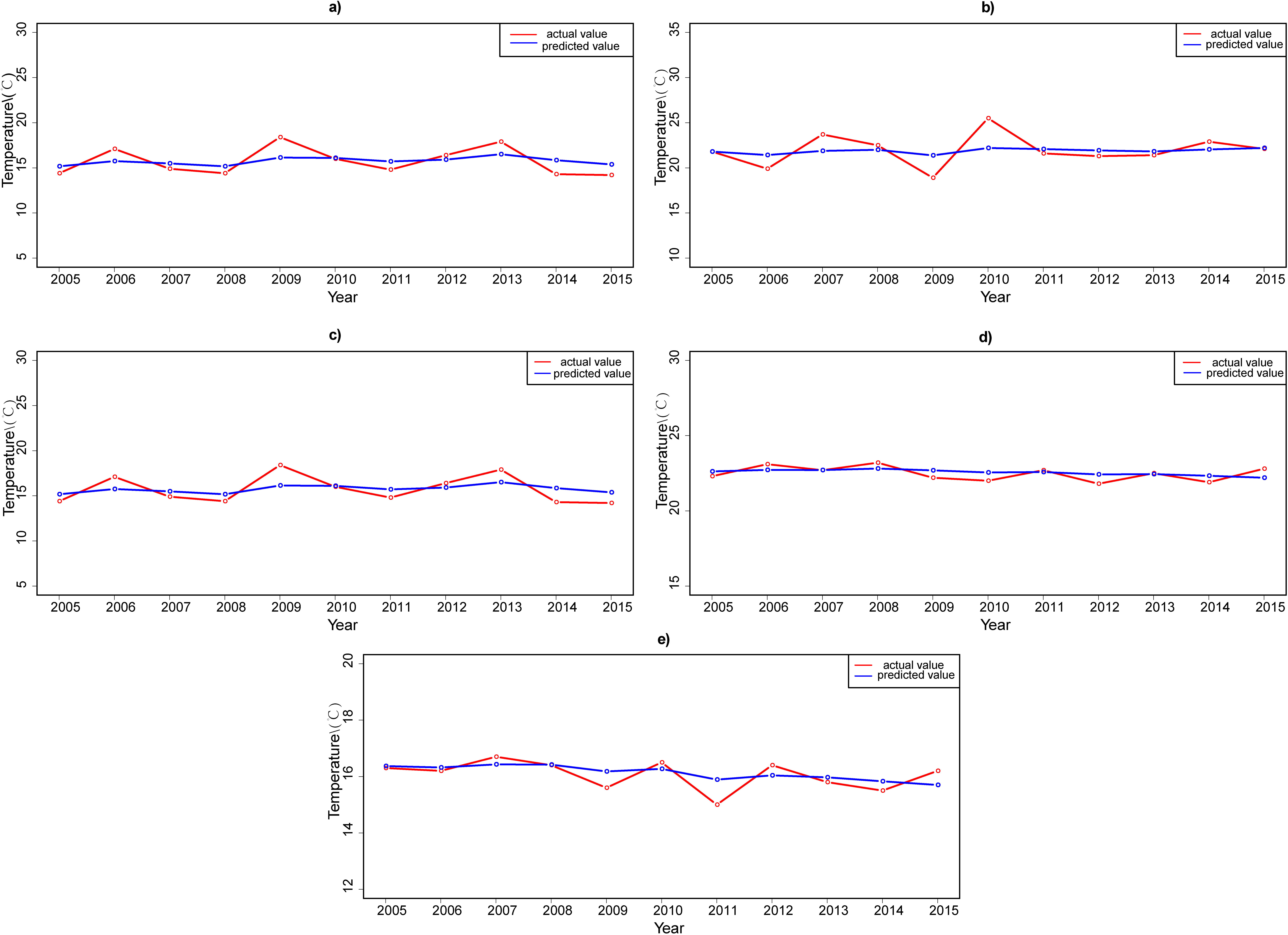

### 3.2 Evaluation classification results based on Euclidean distance

The multidimensional matrix *D* is composed of the actual value of the monthly precipitation, the monthly mean temperature and the sunshine hours of the Harbin pilot for 2005-2014 years and the predicted value of the same meteorological conditions in 2015. The multidimensional matrix *D* consists of only 5-9 months of meteorological data.

Matrix *D*’was obtained after standardizing the column vectors of the multi-dimensional matrix *D*. Then the Euclidean distance between 2015 and 2005-2014 years was obtained according to equation (5) (Fig. 5). The matrix *D*’of Changchun pilot was obtained in the same way, and the Euclidean distance results were shown in Supplemental Fig. 5 online.

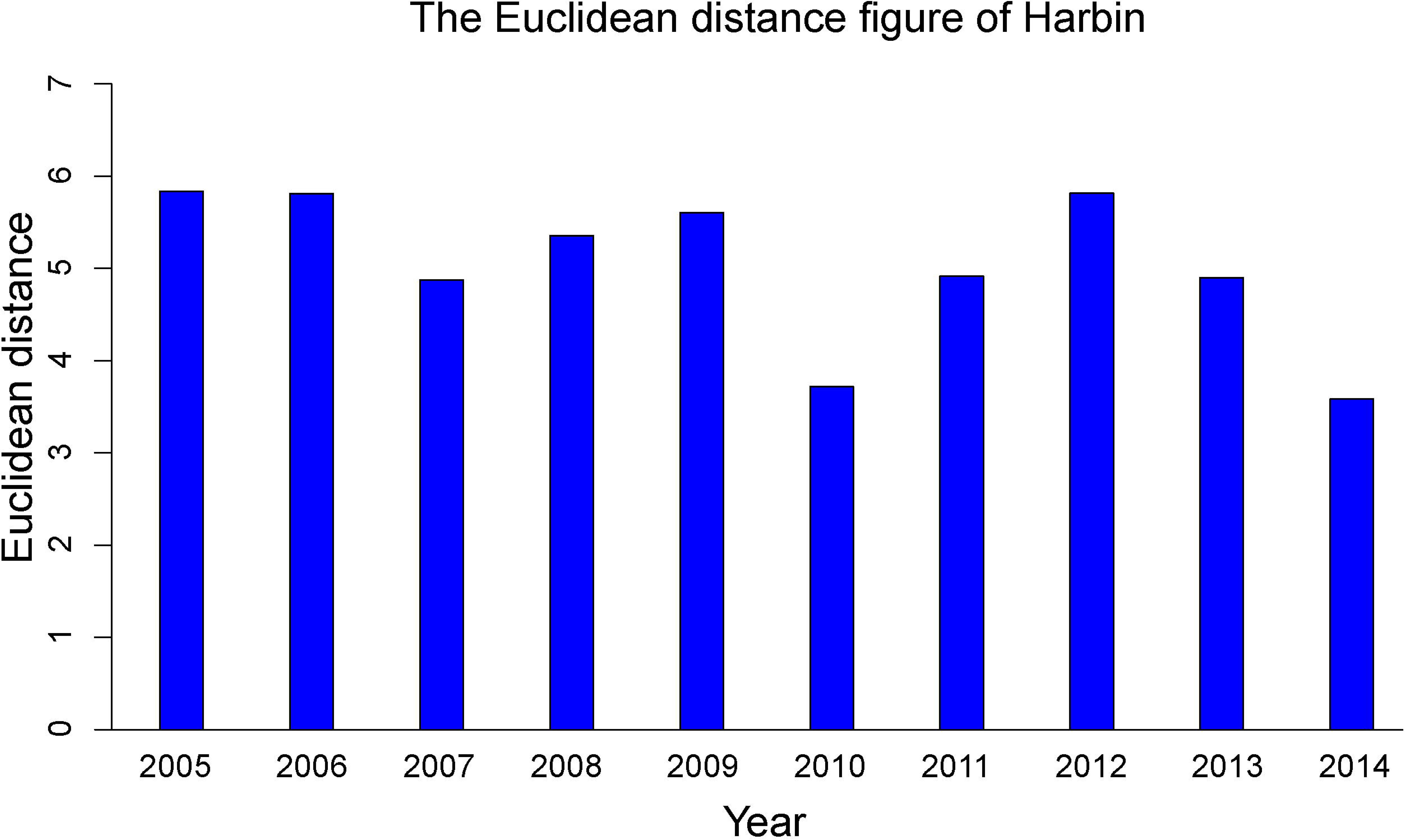

Fig. 5 shows that the shortest Euclidean distance in the Harbin pilot is 2014. Therefore, the meteorological conditions of Harbin pilot in 2014 are similar to the meteorological conditions in 2015, so the varieties planted in 2014 can be recommended to be planted in 2015.

The yield evaluation traits in the experiment were 100-seed-weight and seed weight per plant. Therefore, the actual varieties of soybean in 2014 were sorted by 100-seed-weight and seed weight per plant respectively. Then the threshold α was selected according to the actual needs, and the cultivated varieties in 2015 were selected through the sorting data of the actual soybean varieties in 2014. Supplemental Fig. 5 online shows that the shortest Euclidean distance in the Changchun pilot is 2008. Therefore, the meteorological conditions of Changchun pilot 2008 are similar to that of 2010. The cultivated varieties in 2008 were recommended to be planted in 2010. The yield evaluation traits were also 100-seed-weight and seed weight per plant, and the selection of soybean varieties in Changchun in 2010 will be realized through the same method.

According to 100-seed-weight and seed weight per plant, the actual soybean varieties in 2014 and 2015 in Harbin pilot were sorted, and the actual soybean varieties in 2008 and 2010 in the Changchun pilot were also sorted (Table 3; Table 4). Table 3 and Table 4 enumerate only the varieties required for the experiment, and more sorting varieties refer to the Supplemental Table S7 and Table S8.

**Table 3.**
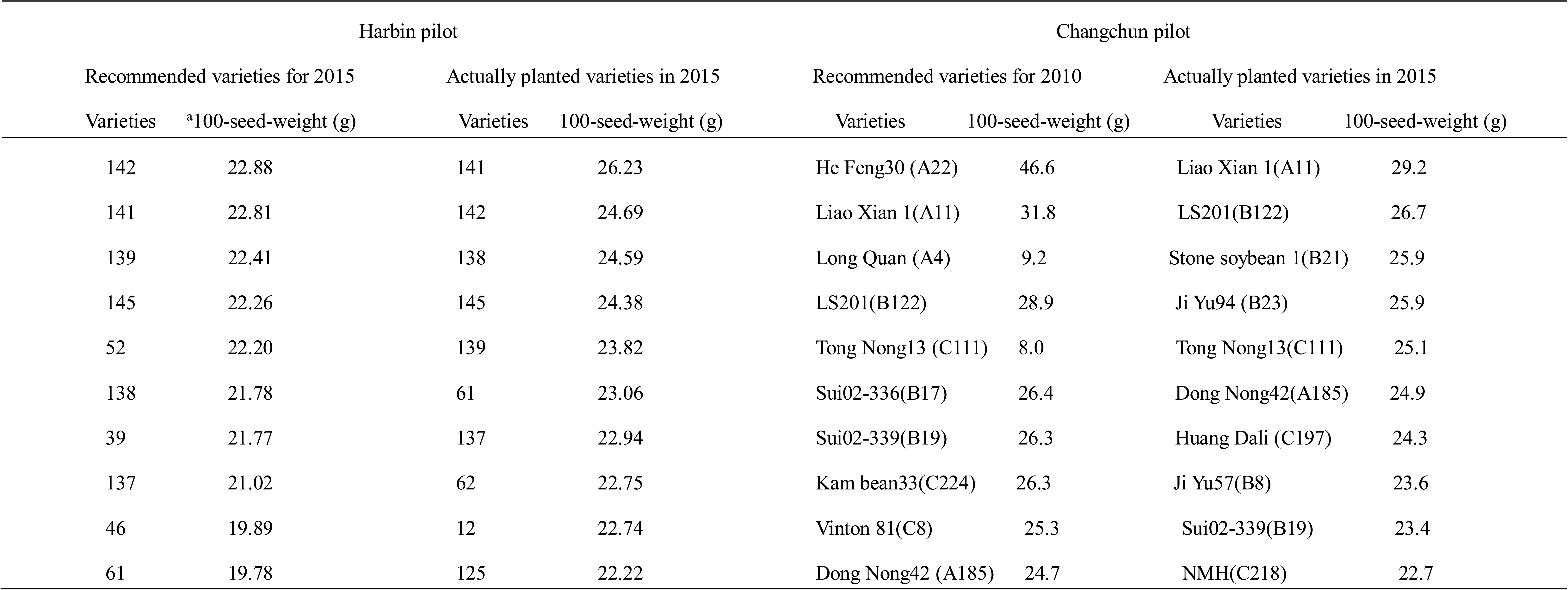
The actual planting varieties of Harbin pilot in 2014 and 2015 and Changchun pilot in 2008 and 2010 were ranked according to 100-seed-weight. 100-seed-weight is an indicator of the size and richness of seeds, which is usually used in large seeds such as corn, soybeans, peanuts, cotton and so on. Expression of varieties: the name of the variety (the community of planting the varieties). For example, He Feng30 (A22) refers to the name of the variety is He Feng30, and the planting community is A22.

**Table 4.**
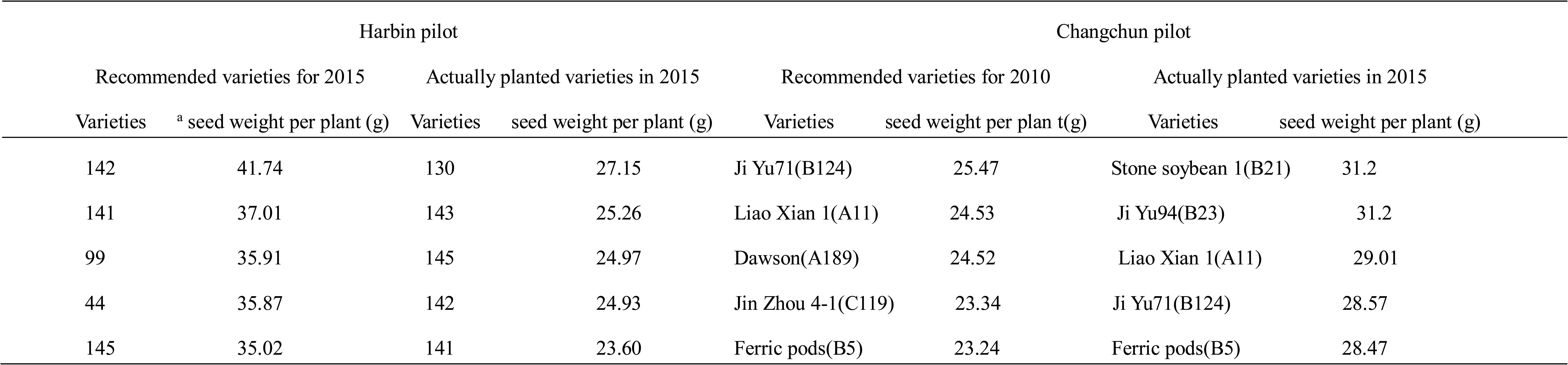
The actual planting varieties of Harbin pilot project in 2014 and 2015 and Changchun in 2008 and 2010 were ranked according to seed weight per plant. Seed weight per plant refers to the total weight of all the seeds harvested from a single plant.

Fig. 6a shows the density distribution of yield evaluation traits (100-seed-weight) of the Harbin pilot in 2010. The density distribution of 100-seed-weight in the 2008 year of Changchun pilot is shown in Supplemental Fig. 6a online. The threshold value was selected according to the actual requirements of different pilots during the selection process. For example, suppose that the threshold α of the Harbin pilot was selected 0.07 to carry out the selection of varieties. Therefore, the selection of varieties was carried out at the high-end area (the right side of Fig. 6a). The total number of varieties in Harbin pilot in 2010 was 149, and the top ten varieties were selected according to the threshold. It was assumed that the threshold α was 0.04 for selection of varieties in Changchun pilot. The total number of varieties in Changchun was 274 in 2008, so the top ten varieties were selected. For 100-seed-weight, The final recommended varieties in the Harbin pilot were 142, 141, 139, 145, 52, 138, 39, 137, 46, 61. The final recommended varieties in the Changchun pilot were He Feng30 (A22), Liao Xian 1(A11), Long Quan (A4), LS201(B122), Tong Nong13 (C111), Sui02-336(B17), Sui02-339(B19), Kam bean 33(C224), Vinton 81(C8), Dong Nong42 (A185) (Table 3).

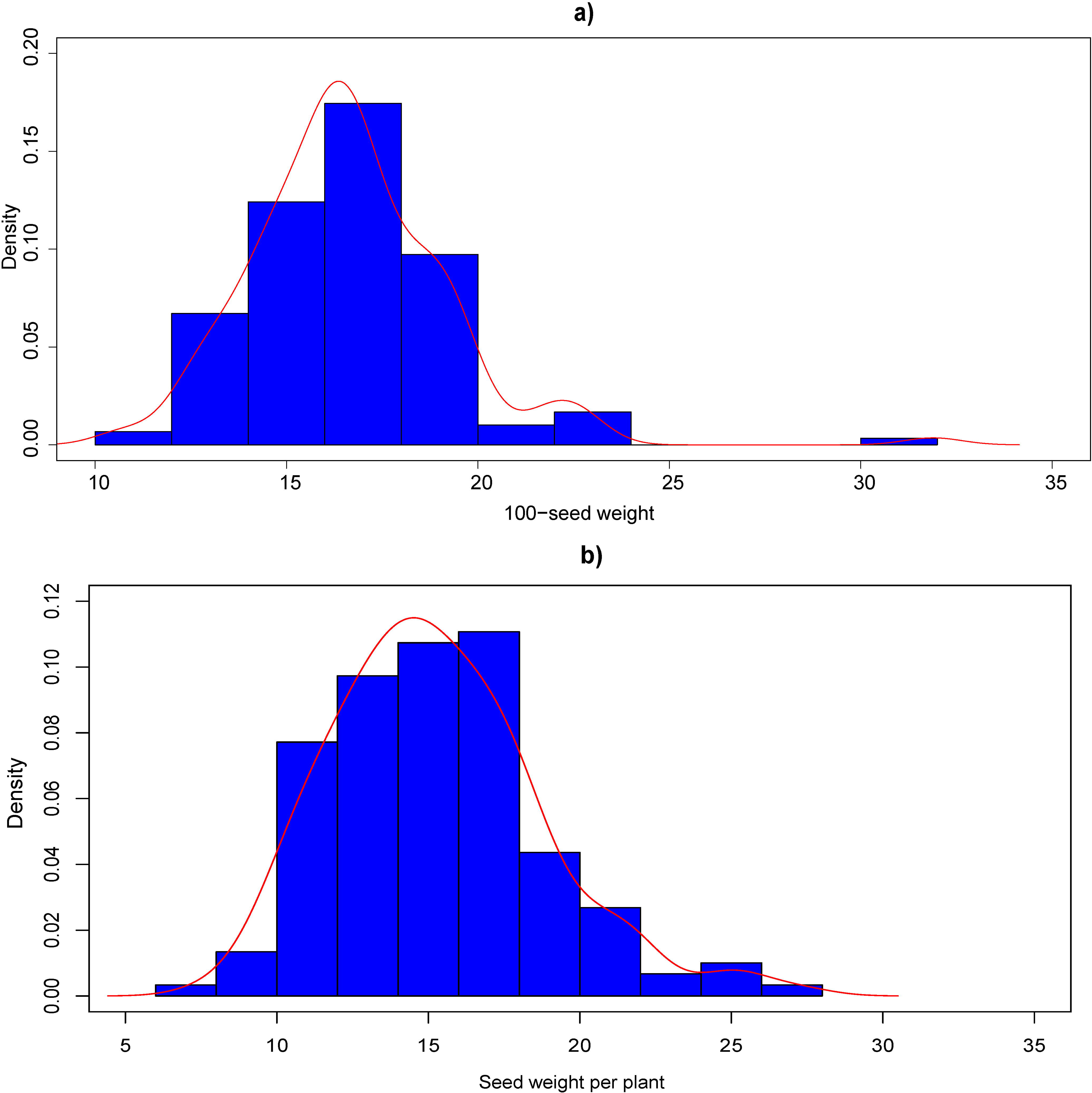

Fig. 6b shows the density distribution of yield evaluation traits (seed weight per plant) of the Harbin pilot in 2010. The density distribution of seed weight per plant in the 2008 year of Changchun pilot is shown in Supplemental Fig. 6b online. For seed weight per plant, the threshold α of the Harbin pilot was selected 0.03 to carry out the selection of varieties. Therefore, the selection of varieties was carried out at the high-end area (the right side of Fig. 6b). The total number of varieties in Harbin pilot in 2010 was 149, and the top five varieties were selected according to the threshold. It was assumed that the threshold α was 0.02 for selection of varieties in Changchun pilot. The total number of varieties in Changchun was 274 in 2008, so the top five varieties were selected. Therefore, the final recommended varieties in the Harbin pilot were 142, 141, 99, 44, 145. The final recommended varieties in the Changchun pilot were Jiyu71(B124), Liao Xian 1(A11), Dawson(A189), Jinzhou 4-1(C119), Ferric pods(B5) (Table 4).

The top ten varieties of 100-seed-weight in 2015 were 141, 142, 138, 145, 139, 61, 137, 62, 12, 125 in Harbin pilot (Table 3). Therefore, it is concluded that there were 7 recommended varieties of soybean in the top ten of the actual ranking in 2015. The top ten varieties of 100-seed-weight in 2010 were Liao Xian 1(A11), LS201(B122), Stone soybean 1(B21), Ji Yu94 (B23), Tong Nong13(C111), Dong Nong42(A185), Huang Dali(C197), Ji Yu57(B8), Sui02-339(B19), NMH(C218) in Changchun pilot (Table 3). It was concluded that there were five recommended varieties of soybean in the top ten of the actual ranking in 2010. Therefore, according to the above analysis ratio, the recommended varieties in Harbin pilot and Changchun pilot could have the possibility of 70% and 50% above, respectively, to provide a prediction for our selection.

The top five varieties of seed weight per plant in 2015 were130, 143, 145, 142, 141 in Harbin pilot. In Changchun pilot, the top five varieties in 2010 were Stone soybean 1(B21), Ji Yu94(B23), Liao Xian 1(A11), Ji Yu71(B124), Ferric pods(B5) (Table 4). Therefore, it was concluded that the first five soybean varieties in Harbin pilot in 2015 included three recommended varieties, and the result of Changchun pilot was the same as that of it. So as for the seed weight per plant, the possibility of more than 50% of the recommended varieties in Harbin pilot and Changchun pilot was to provide a pre-judgement for the selection of cultivated varieties. The above analysis is of vital importance to the selection of new varieties for selecting new varieties and increasing yield.

## 4 Discussion

In the study of varieties selection of crops, the multi-objective selection function was established by the multi-objective decision technology which combines the analytic hierarchy process and the utility function method^40^. Liao and Guan completed the combination of computer information technology and agronomy, so the experts’ experience knowledge and the methods of solving problem were inherited and solidified of rapeseed, which laid the foundation for the standardization and informatization of rapeseed cultivation and management^41^. The methods recommended in the above literatures mostly relied on experiential knowledge rules in knowledge base and a large number of regional experiments to achieve the selection of planting varieties, and the effect of meteorological conditions on crop growth was not considered in the process of seed selection.

Though the paper used knowledge engineering and system modeling methods, the dynamic relationship between varieties selection and varieties, ecological environment and farming system was summarized, and a quantitative knowledge model for suitable varieties selection was established^42^. However, there are few articles on yield, disease resistance and phenotypic quality index, so the selection methods based on yield traits is still worth studying.

The meteorological prediction model based on phase space reconstruction and exponential smoothing method can predict meteorological conditions accurately and provide accurate data support for the selection of soybean varieties. However, some meteorological conditions have a variety of time scales, which makes meteorological conditions pretty complicated. If only one prediction model is used in the trial process, the complexity of the time series itself will lead to the inaccuracy of the prediction results. Therefore, if we want to obtain accurate prediction results, we will choose multiple time series prediction models according to the experiment requirements, such as the combination of phase space reconstruction and neural network.

Although the prediction method of phase space reconstruction and based on exponential smoothing in long term prediction is not able to make full use of multi-scale characteristics and regularities of the variation of the meteorological conditions. Chaotic space theory can make better use of the information contained in the time series to solve the problems that are difficult to solve in the prediction model of general nonlinear meteorological conditions in the short-term prediction, which can predict the meteorological conditions more accurately and meet the requirement of this strategy.

The meteorological data of monthly precipitation, monthly mean temperature and monthly sunshine hours in Harbin during the past 2005-2015 years were selected as meteorological materials for Harbin pilot in this paper. However, the Changchun pilot only used meteorological data from 2008 to 2010. In Harbin pilot, 2013-2015 years of meteorological data can also be used for the application of the strategy, and the concrete experiment steps are as follows.

**Step 1:** The meteorological data from 2013 to 2014 were selected as training samples, and the meteorological data in 2015 were testing samples. Meteorological data of 2015 were obtained by using the meteorological prediction model in the text (Table 5).

**Table 5.**
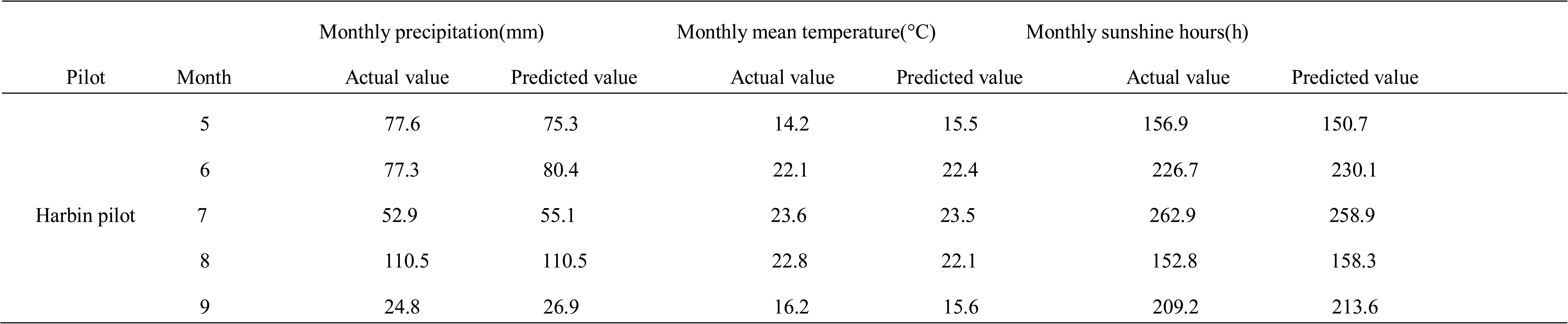
Comparison of forecast data and actual data of meteorological conditions in Harbin in 2015. In this form, the meteorological data in 2013-2014 years are used as experimental materials to predict the meteorological conditions in 2015 through the meteorological prediction model in the text.

**Step 2:** According to equation (1), the meteorological data of 2013-2015 years (the meteorological data of 2015 as the prediction data) were standardized. Then, according to the evaluation classification model based on Euclidean distance and equation (5), the Euclidean distance between 2015 and 2013 was 4.78, and the Euclidean distance between 2015 and 2014 was 4.80. Therefore, the meteorological conditions in 2015 were similar to that in 2013, so the soybean varieties in 2013 could be the preferred planting varieties in 2015. The density distribution of 100-seed-weight of soybean varieties in Harbin pilot in 2013 was shown in Fig. 7.

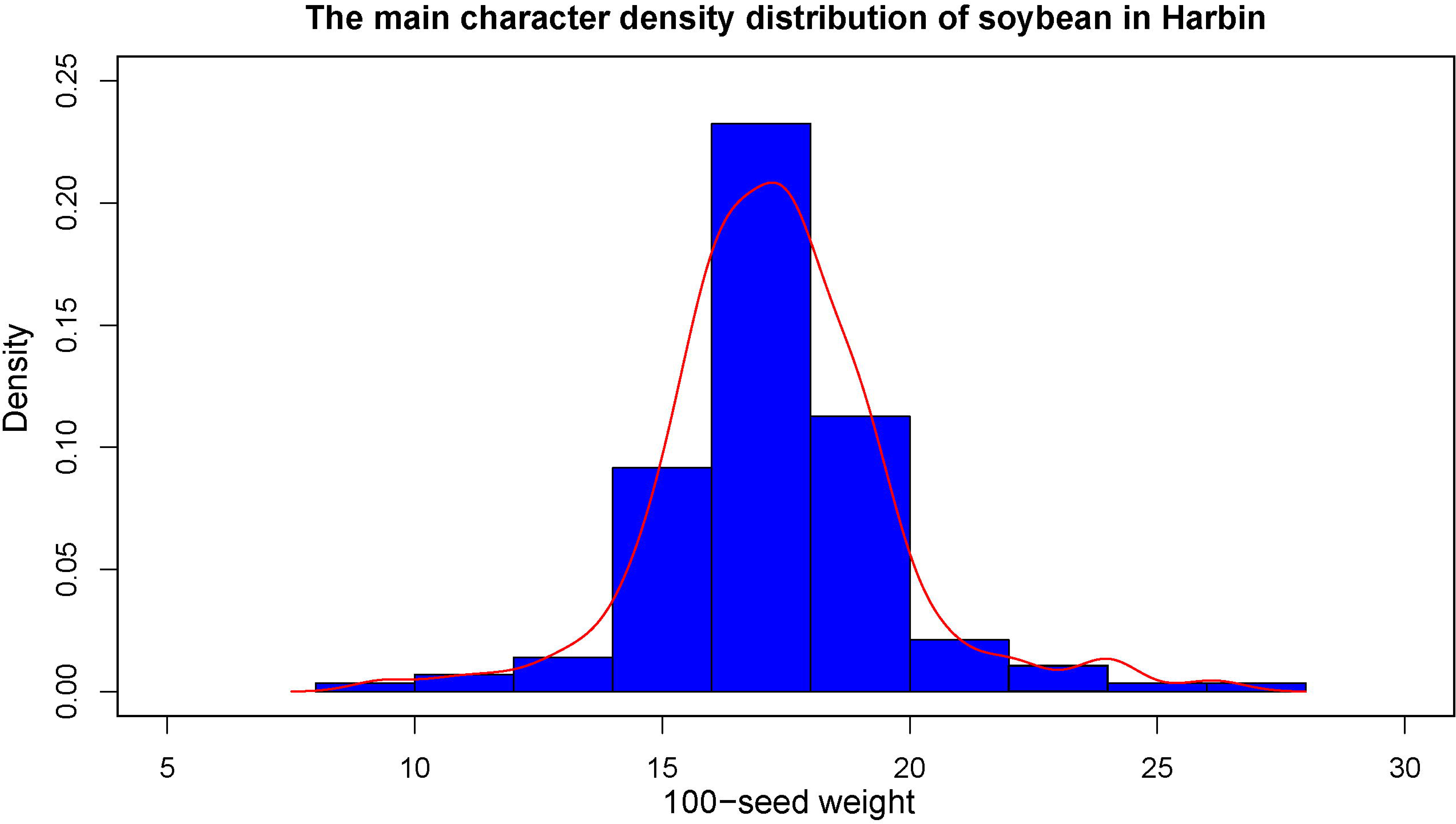

In this experiment, the threshold α was also selected to be 0.07, and the selection of the varieties could be made in the high-end area (the right side of Fig. 7). According to the Supplemental Table S7, the top ten varieties of 100-seed-weight in 2013 were: 142, 145, 138, 68, 52, 141, 39, 21, 139, 127. In 2015, the top ten varieties of actual soybean were 141, 142, 138, 145, 139, 61, 137, 62, 12, 125. It is easy to conclude that there were five kinds of recommended planting varieties in 2015. The method was to use the meteorological data from 2013 to 2015 for the application of this model. The results showed that even three years of meteorological data also can provide more than 50% possibilities for planting varieties in the coming year, which was also of great practical significance for improving soybean yield. This part only studied the 100-seed-weight, and the evaluation method of seed weight per plant is the same as that of the text.

In this paper, the selection model of single target and single location was discussed, and the model of single target and multi-location, multi-target and single location, and multi-target and multiple locations.

As for the selection model of single target and multi-location, the locations can be recorded as: *P*_1_, *P*_2_, …, *P*_*m*_. The meteorological years to be studied can be recorded as *C*_1_, *C*_2_, …, *C*_*s*_. Therefore, the environmental factors changed from the previous *S* to the *P* * *S*. Then it will be selected according to the model of single target and single location.

As for the selection model of multi-target and single location, the following method can be used. Suppose there are *n* species and *m* traits:

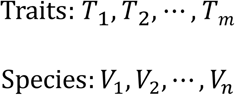

For example, *s*(*s* ≤ *n*) species are selected for the target trait *T*_*i*_(*i* ∈ {1, *m*}). The varieties are expressed as follows:

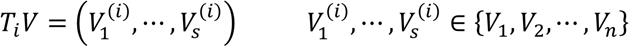

The soybean varieties are sorted by the size of the main target trait *T*_*i*_(*i* ∈ {1, *m*}):

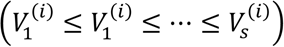

For different traits *T*_1_, *T*_2_, …, *T*_*m*_, the sequence vectors of the selected varieties are as follows:

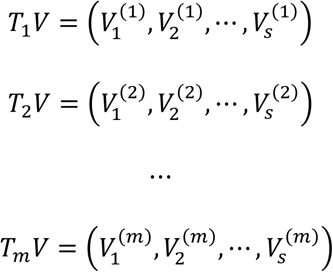

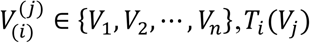 represents the value of the *i* trait of the *j* variety, *i* ∈ {1, m}, *j* ∈ {1, n}.

In the selection of multi-target, the required yield main trait can be selected according to the single target selection strategy, and the minimum value of the remaining traits is then defined according to the actual needs. Suppose the minimum value of the yield main character is set to *∂*, and there is a variety of 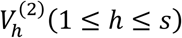 in *T*_2_*V*, which does not appear in *T*_1_*V*. Then the yield main character value of 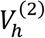 is required. If the yield main character value of the 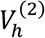 is higher than *∂*, the *T*_1_*V* is reordered after the 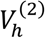 is inserted into the *T*_1_*V* according to the value of the yield main trait of 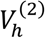. If the yield main character value of 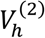 is lower than *∂*, it will be discarded directly. According to the above methods, the selection of multi-target traits can be achieved.

As for the selection model of multi-target and multiple locations, the environmental factors of multi-location can be determined according to the model of single target and multi-location. The selection of multi-target can be determined according to the selection model of multi-target and single location.

The classification model of crop cultivated varieties can accurately classify the varieties according to Euclidean distance. The method in the text is carried out after the standardization of the column vector of the meteorological condition matrix *D*. The method discussed here is to calculate the Euclidean distance between the three meteorological conditions respectively.

Because the Euclidean distance is calculated by the same meteorological conditions, there is no intrinsic difference between the same weather conditions, and standardization is not used in the model. The specific model is as follows:

Suppose the meteorological data matrix for *n* years is *A*.

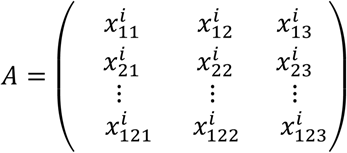

The *i*(1 ≤ *i* ≤ *n*) in the matrix means the number of years of the observed value. The three columns in the matrix represent monthly precipitation, monthly mean temperature and monthly sunshine hours respectively. Based on the phase space reconstruction and exponential smoothing prediction model, the matrix *B* that consists of the predicted data of monthly precipitation, monthly mean temperature and monthly sunshine hours of *n* + 1 year can be obtained.

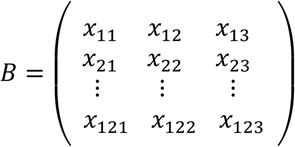

The idea of the evaluation classification model is to get the Euclidean distance *d* of each column vector of the prediction matrix *B* and the observation matrix *A* separately. The approximate degree of the prediction matrix *B* and the observation matrix *A* is judged by the size of the Euclidean distance. The smaller the Euclidean distance, the greater the degree of similarity, and then *Y*_1_, *Y*_2_ and *Y*_3_, which correspond to the shortest Euclidean distances of the three meteorological conditions respectively, are selected. The Euclidean distance can be calculated according to the monthly precipitation calculation formula equation (8), monthly mean temperature formula equation (9) and monthly sunshine hours calculation formula equation (10).

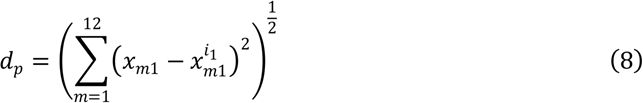

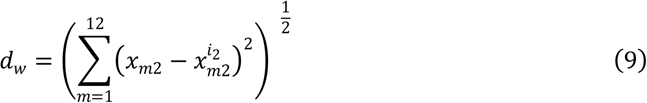

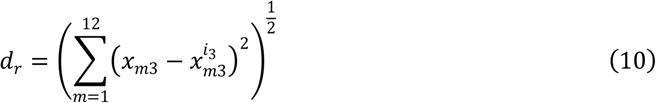

Taking the Harbin pilot as an example, this method can be used to obtain the Euclidean distance between the observation matrix *A* and the prediction matrix *B* in Harbin pilot (Fig. 8), and the results of the Changchun pilot are shown in Supplemental Fig. 7 online.

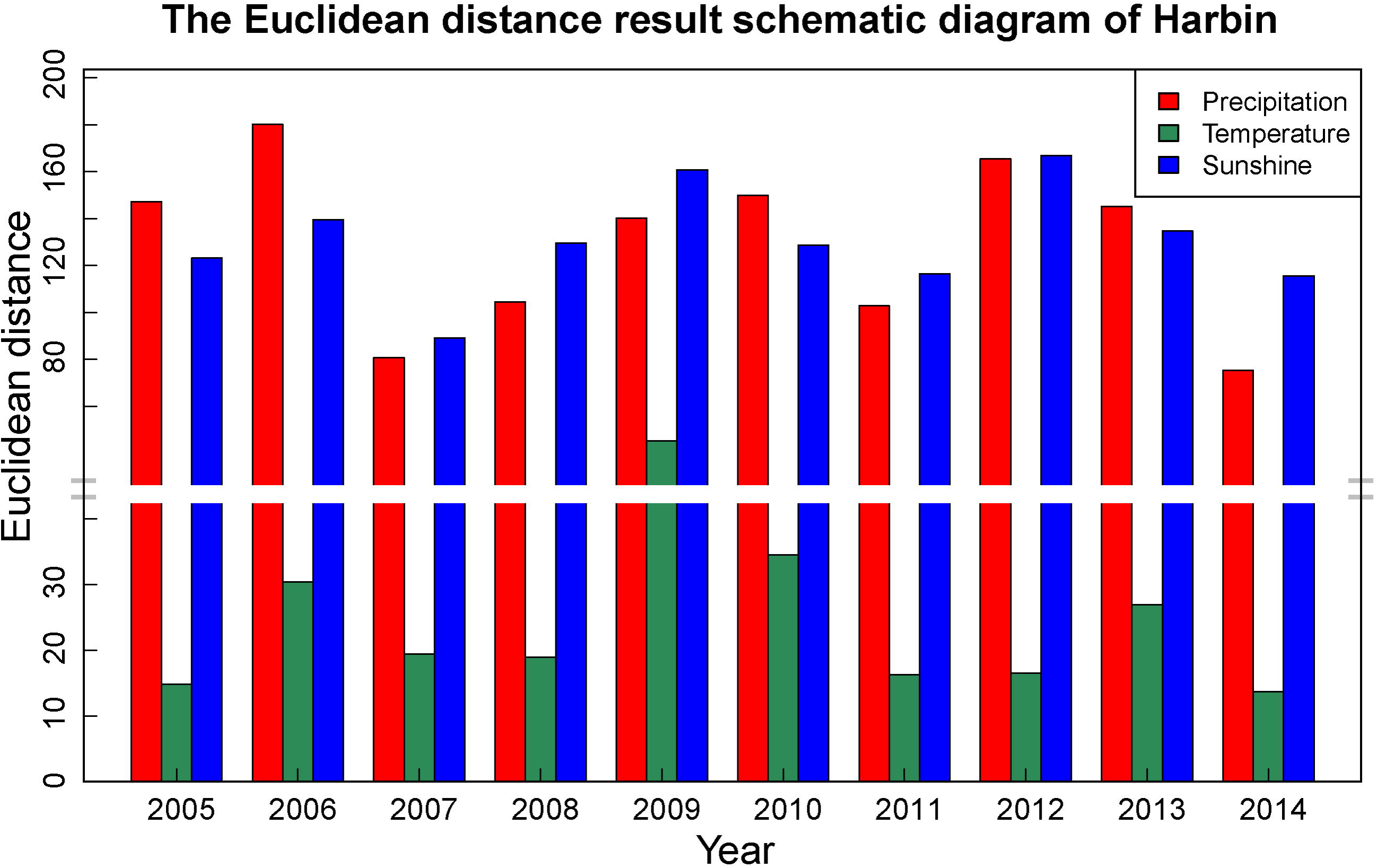

In this method, the 100-seed-weight and seed weight per plant can still be used as yield evaluation traits. Here, only 100-seed-weight is used as an example. Fig. 8 shows that the monthly precipitation and monthly average temperature of the Harbin pilot in 2015 are similar to those in 2014, and the weather condition of the monthly sunshine hours is similar to that in 2007. In 2007 and 2014, the data of the top ten varieties of soybean 100-seed-weight were shown in Table 6. Supplemental Fig. 7 online shows that the monthly precipitation, monthly average temperature and sunshine hours of Changchun in 2010 are similar to those of 2008, and the selection of planting varieties can be realized by the same method.

**Table 6.**
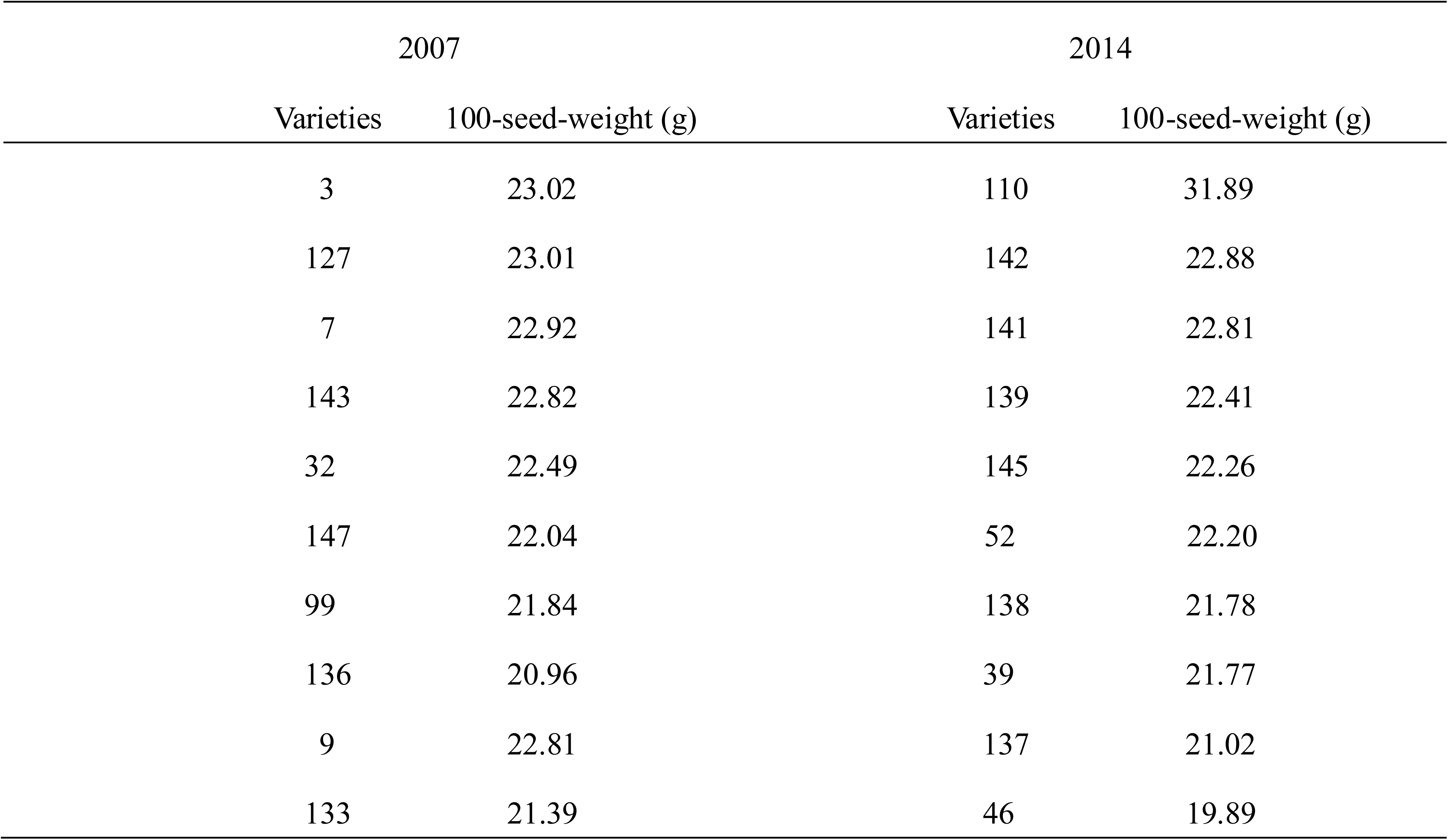
The ranking of 100-seed-weight of soybean varieties in 2007 and 2014 in Harbin pilot. Fig. 8 shows that the monthly precipitation and monthly average temperature of the Harbin pilot in 2015 are similar to those in 2014, and the weather condition of the monthly sunshine hours is similar to that in 2007.

The weight of different meteorological factors that affect the growth of soybean needs to be considered when selecting the recommended planting varieties by this method. Because of the different areas of planting crops, the weight of meteorological factors will be different. Therefore, the weight of the three meteorological conditions should be allocated according to the meteorological conditions of the planting area, and then the selection of crop cultivated varieties should be carried out. The specific steps are as follows:

**Step 1:** It is assumed that the total number of crop cultivated varieties planted in *Y*_1_, *Y*_2_ and *Y*_3_ is A, B and C respectively.

**Step 2:** The weight of precipitation, average temperature and sunshine hours is α_1_, α_2_, α_3_, respectively, according to the meteorological conditions of the actual planting area.

**Step 3:** A, B and C are sorted according to the yield evaluation traits, and the number of varieties will be selected according to different weights. Therefore, the total number of varieties selected in *Y*_1_, *Y*_2_ and *Y*_3_ is A · *α*_1_, B · *α*_2_ and C · *α*_3_ respectively.

**Step 4:** The final set of recommended planting variety is *P* = A · α_1_ ∩ B · α_2_ ∩ C · α_1_.

## 5 Conclusion

In this paper, a new strategy based on meteorological driving for classification and selection of cultivated varieties is presented. The main purpose is to select cultivated varieties based on the prediction of meteorological conditions so as to increase crop yield. The strategy realizes the combination of the meteorological prediction model based on phase space reconstruction and exponential smoothing and the variety evaluation classification model based on Euclidean distance. In the experiment, the strategy accurately grasped the interactive relationship between soybean yield and meteorological conditions, which provided a scientific basis for soybean to make full use of meteorological resources, avoid disadvantages and increase yield. Not only can this strategy complete the selection of soybean varieties, but also it can be extended to other crops’ seed selection, so it has broad application prospects for the selection of crop cultivated varieties in the agricultural field.

## Author Contributions

Rongsheng Zhu, Kai Sun put forward the strategy and wrote this paper. Xuehui Yan supplied the meteorological data weather data. Zhaoming Qi and Chunming Yang supplied the yield data of soybean. Dawei Xin, Hongwei Jiang, Zhanguo Zhang, Yang Li, Zhenbang Hu designed the experiment and modified this paper. Rongsheng Zhu, Kai Sun and Qingshan Chen modified this paper.

## Additional Information

### Data availability statement

The data involved in this paper are open.

