## Supplementary figures and images for "A new strategy for selection of crop cultivated varieties based on yield factors driving and meteorological prediction"

### Supplemental Fig. S1

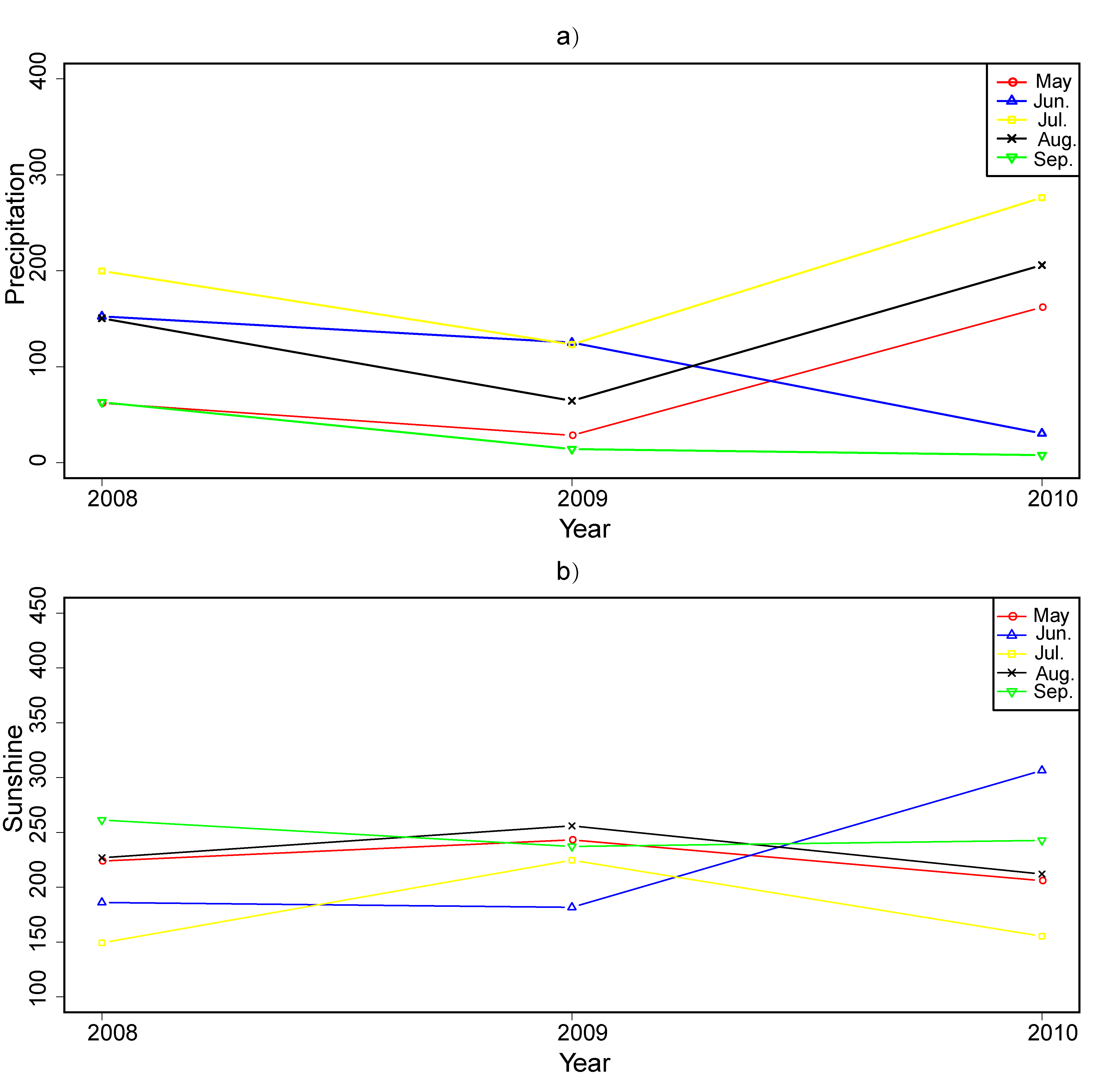

### Supplemental Fig. S2

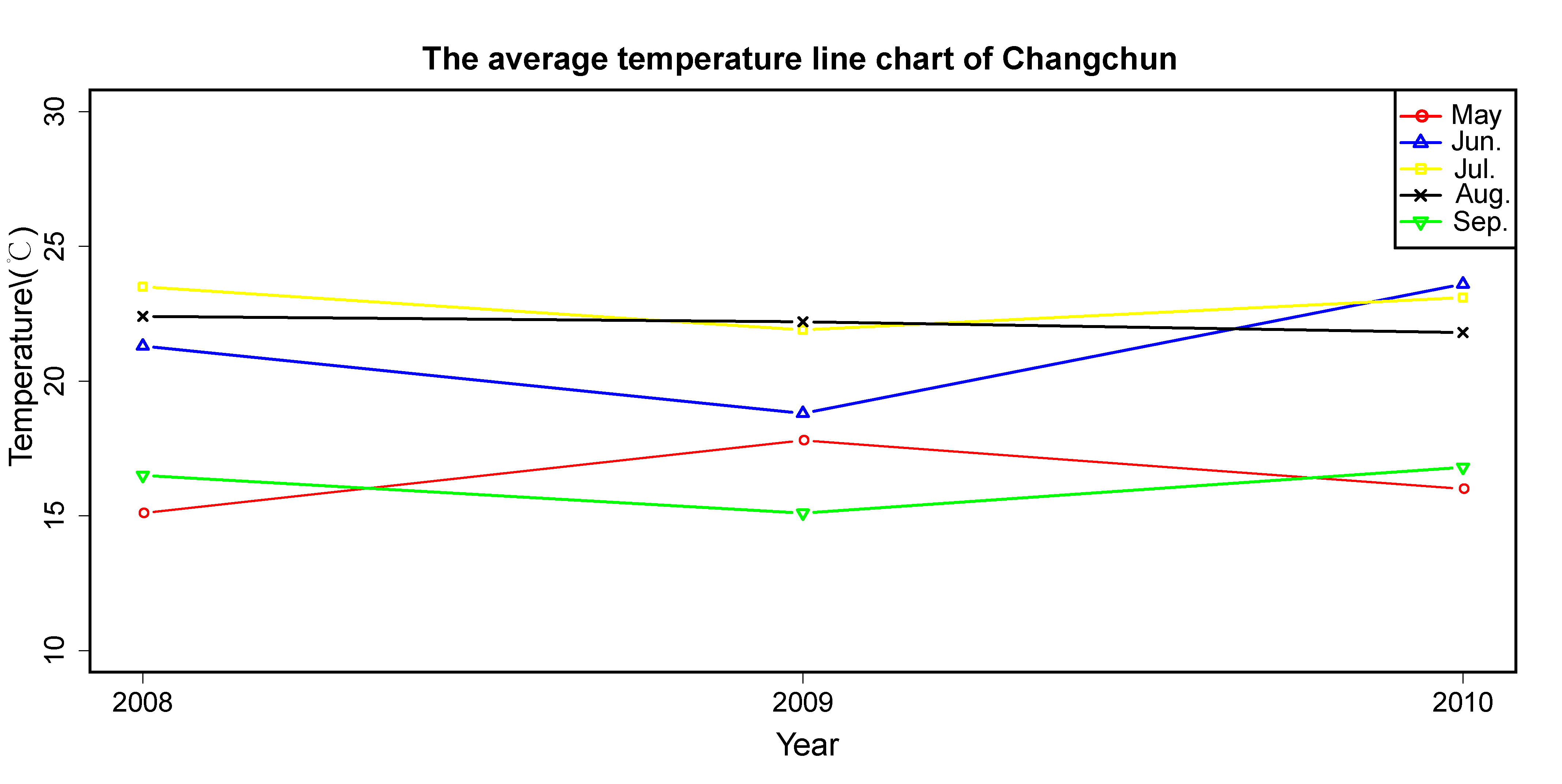

### Supplemental Fig. S3

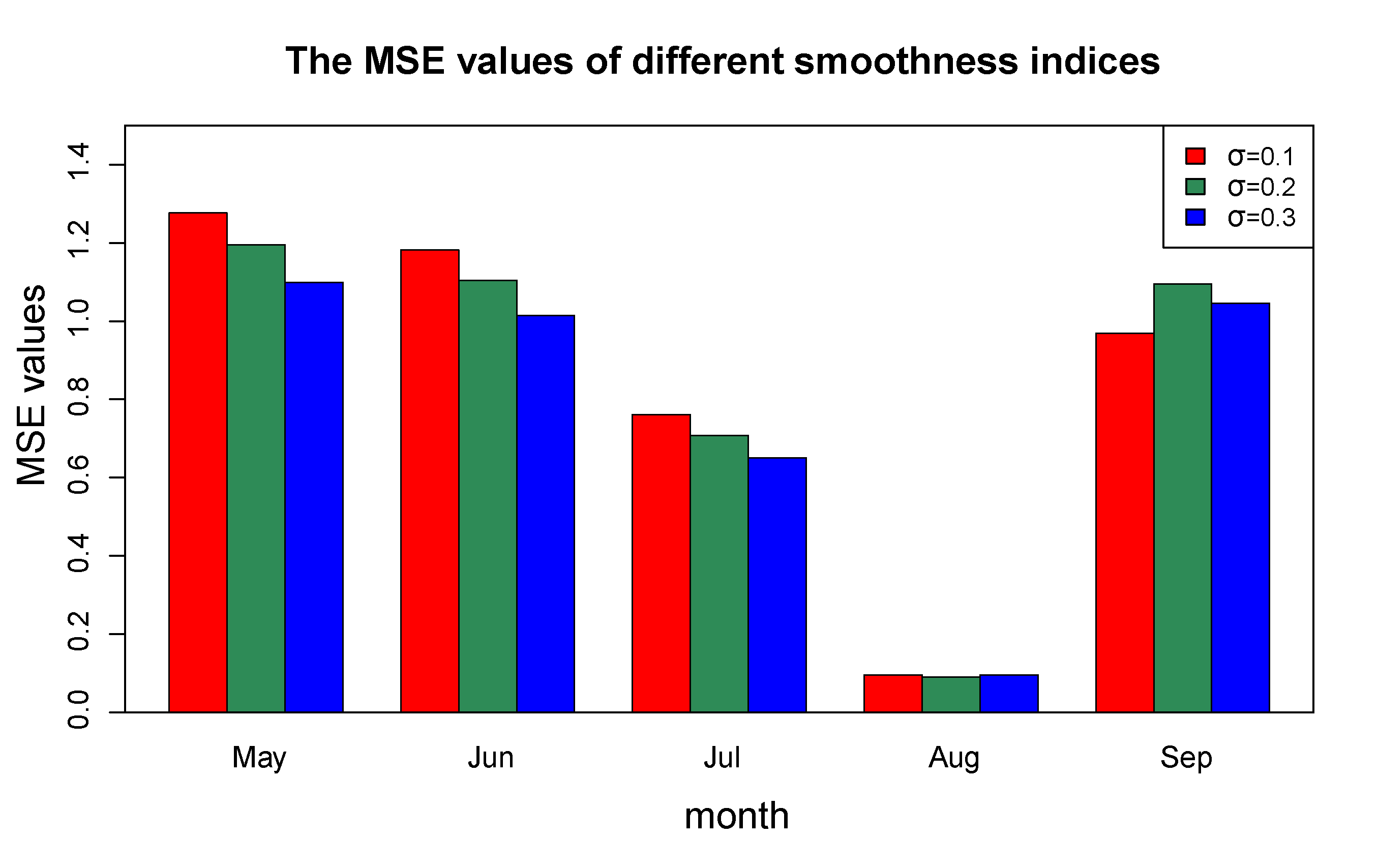

### Supplemental Fig. S4

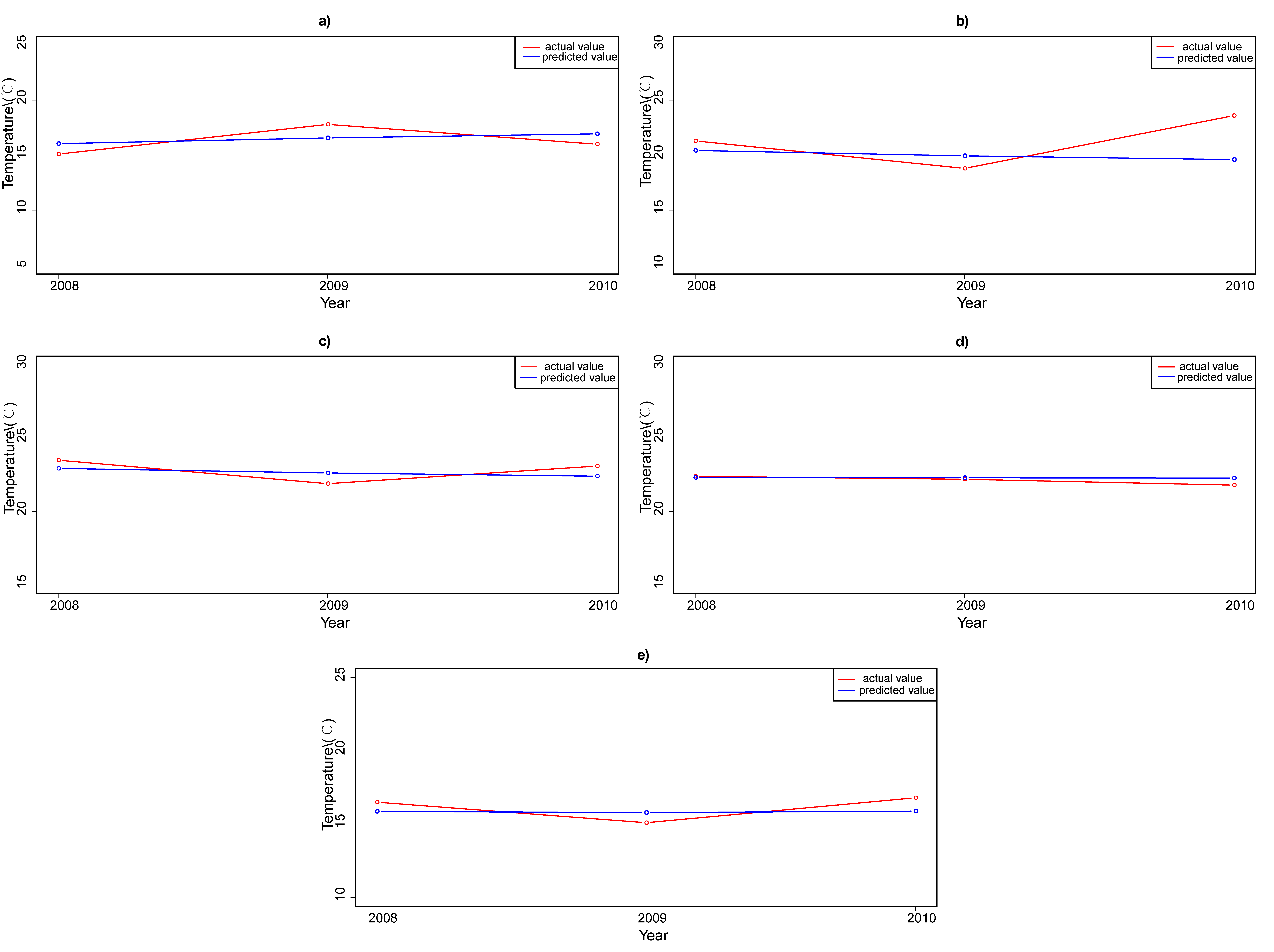

### Supplemental Fig. S5

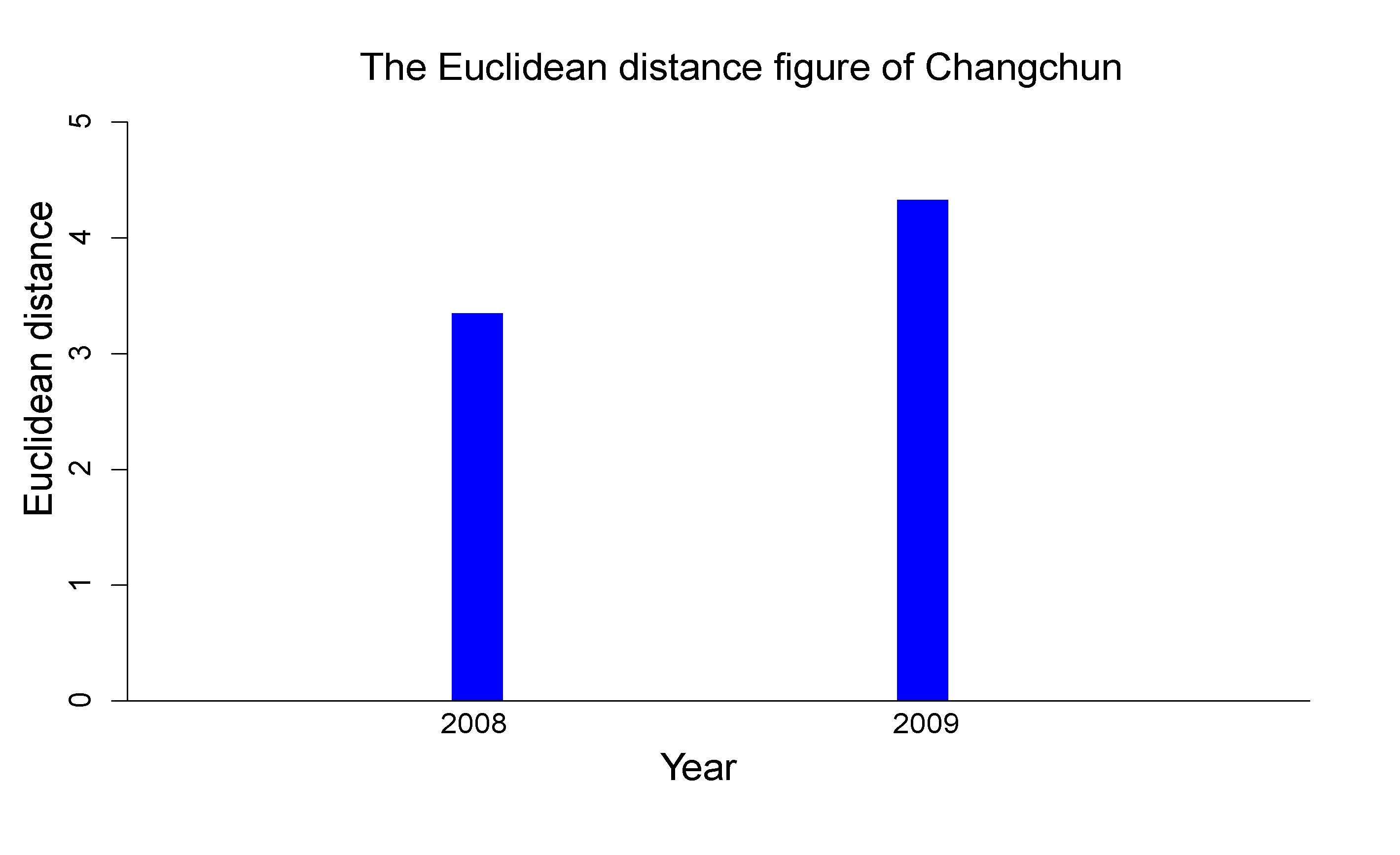

### Supplemental Fig. S6

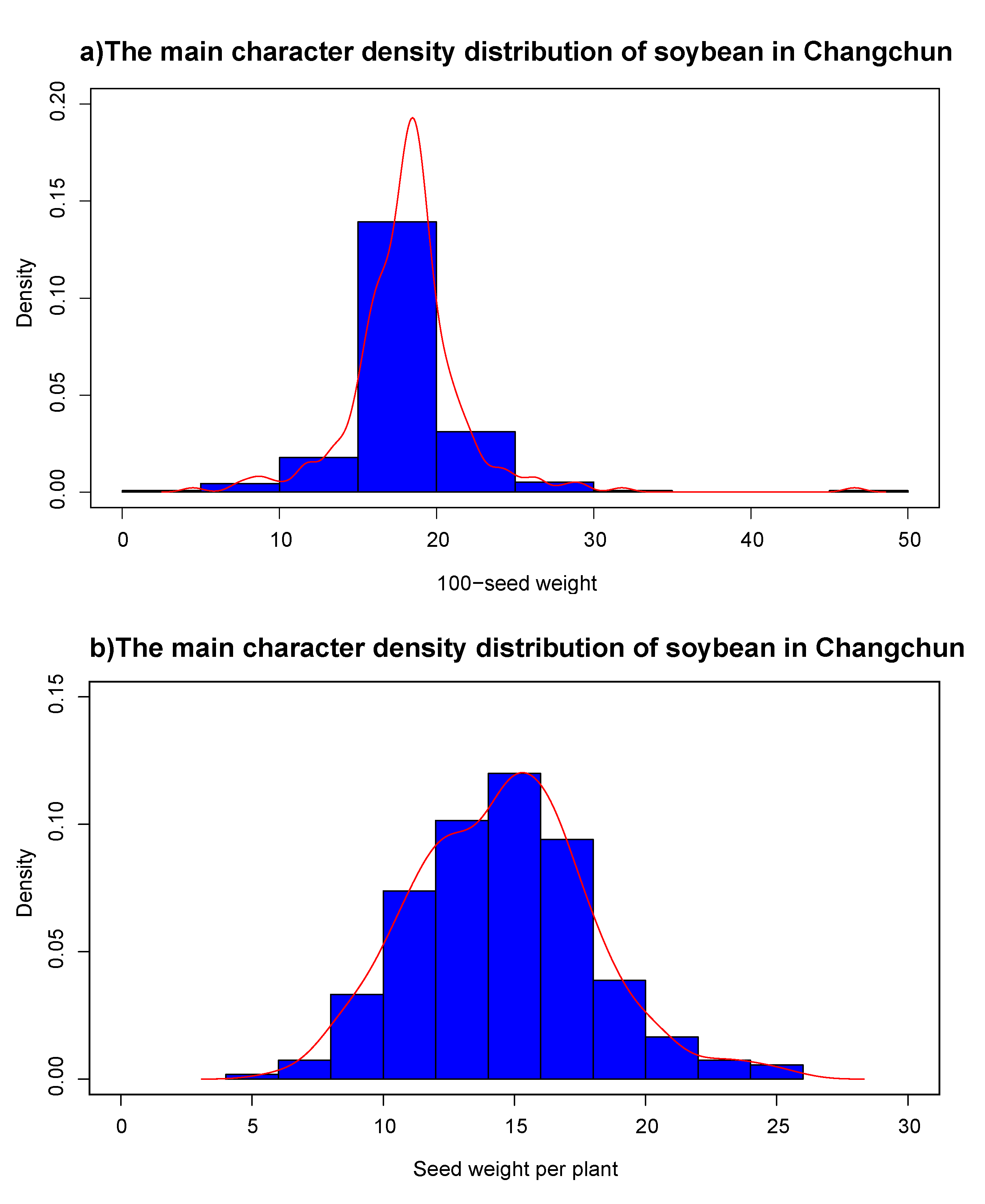

### Supplemental Fig. S7

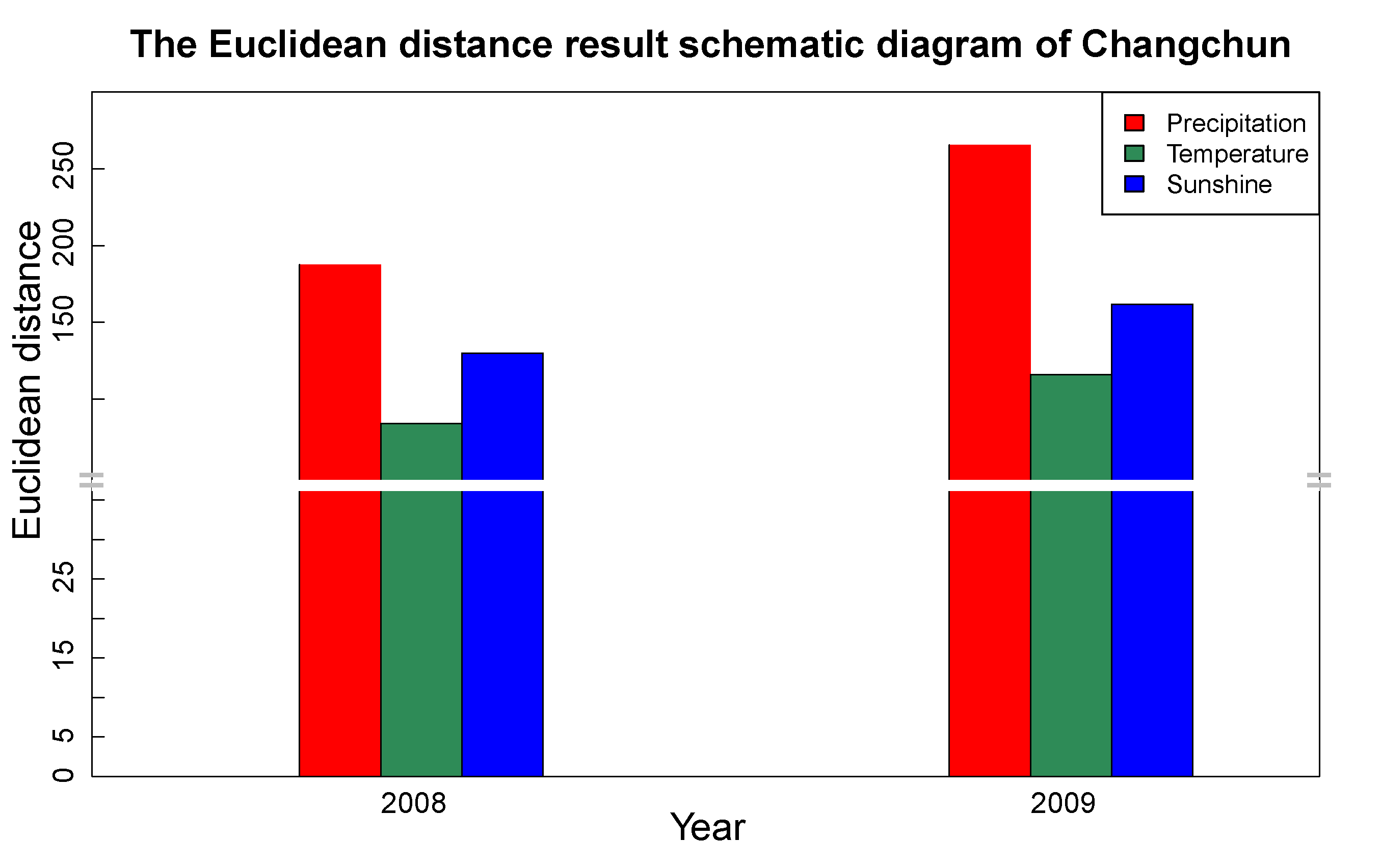
